# Human senataxin is a bona fide R-loop resolving enzyme and transcription termination factor

**DOI:** 10.1101/2022.08.25.505353

**Authors:** Zdenka Hasanova, Veronika Klapstova, Odil Porrua, Richard Stefl, Marek Sebesta

## Abstract

Prolonged pausing of the transcription machinery may lead to the formation of three-stranded nucleic acid structures, called R-loops, typically resulting from the annealing of the nascent RNA with the template DNA. Unscheduled persistence of R-loops and RNA polymerases may interfere with transcription itself and other essential processes such as DNA replication and repair. Senataxin (SETX) is a putative helicase, mutated in two neurodegenerative disorders, which has been implicated in the control of R-loop accumulation and in transcription termination. However, understanding the precise role of SETX in these processes has been precluded by the absence of a direct characterisation of SETX biochemical activities. Here, we purify and characterise the helicase domain of SETX in parallel with its yeast orthologue, Sen1. Importantly, we show that SETX is a bona fide helicase with the ability to resolve R-loops. Furthermore, SETX has retained the transcription termination activity of Sen1 but functions in a species-specific manner. Finally, subsequent characterisation of two SETX variants harbouring disease-associated mutations shed light into the effect of such mutations on SETX folding and biochemical properties. Altogether, these results broaden our understanding of SETX function in gene expression and the maintenance of genome integrity and provide clues to elucidate the molecular basis of SETX-associated neurodegenerative diseases.

## INTRODUCTION

Essential cellular processes such as replication, recombination, and DNA repair occur alongside transcription on DNA. During transcription, RNA polymerase II (RNAPII) pauses at discrete sites, which in some instances is required for proper regulation of gene expression (1, 2). Prolonged pausing and/or stalling of the transcribing RNAPII occurring either for regulatory purposes or under pathological conditions may impinge upon genomic functions by creating an obstacle for DNA-associated machineries (1, 3). RNAPII pausing favours the appearance of R-loops, which are nucleic-acid structures formed when the nascent RNA anneals back to the complementary template DNA strand, thereby leaving the antiparallel non-template DNA strand unpaired. If they remain unprocessed, R-loops may present a source of genome instability. On one side the exposed single-strand DNA (ssDNA) is vulnerable to cleavage by endonucleases and oxidative damage (4, 5). On the other side, R-loops may represent a block for DNA replication and repair (6–11). R-loops are indeed important contributors of transcription-replication conflicts (TRCs), which occur when the replication fork encounters a transcriptional block. Cells possess multiple factors at the interface of the replication and transcription machineries, to overcome TRCs, some of which are involved in the direct removal of R-loops and/or dislodging of RNAPIIs to enable replication fork restart (1, 12–23).

Not only R-loops but also stalled RNAPIIs itself may act as a barrier for transcription, DNA replication, and DNA repair (1). Timely termination of transcription and removal of RNAPIIs and R-loops is, therefore, critical to maintain genome stability, successfully transmit genetic information across generations, and to timely and accurately execute the transcriptional programme.

Aside from their potentially harmful nature, R-loops also play a role in various important physiological processes (21, 24) such as the regulation of gene expression (25–27), the regulation of homologous recombination during DNA repair and immunoglobulin class-switch, and the maintenance of telomeric and centromeric regions (7–9, 11, 28–32). Therefore, R-loop levels need to be tightly regulated to ensure a proper balance between its physiological and pathological roles.

Accumulation of detrimental R-loops may be prevented by mechanisms that avoid the formation of R-loops, for instance, by proper processing of pre-mRNAs, handling of topological changes, and chromatin remodelling during transcription (5, 33–37). Efficient transcription termination, which leads to the release of RNAPII and the RNA transcript from DNA, may also decrease the likelihood of formation of R-loops by reducing the residency time of the RNA in the proximity to the template DNA (38, 39). Once R-loops are formed, additional factors, including nucleases such as RNase H and RNA:DNA helicases, operate to directly remove them (4, 7, 13, 30, 38–47).

A key actor in the control of R-loop accumulation in humans is the putative RNA:DNA helicase, senataxin (SETX), which belongs to the superfamily 1B (SF1B) class of helicases. SETX is homologous to the well-characterised yeast protein Sen1, which plays a major role in transcription termination and the maintenance of genome stability. Detailed biochemical characterisation of Sen1 demonstrated that it is a helicase with the ability to translocate along single-strand DNA and RNA in the 5′ to 3′ direction and to unwind various nucleic acid duplexes (48–50). When Sen1 translocates on the nascent RNA and encounters the transcribing RNAPII, it induces the dissociation of the elongation complex, thereby releasing the RNAPII and the transcript from the DNA (48, 51). During S-phase, Sen1 associates with the replication fork (52, 53) and uses its transcription termination activity to remove RNAPIIs and associated R-loops from TRC sites to aid fork progression (54–56).

Akin to Sen1, SETX has been shown to prevent the accumulation of R-loops at sites of TRCs and/or DNA damage (5, 6, 57–59). Based on the strong conservation between SETX and Sen1 and the demonstrated capacity of the yeast counterpart to unwind RNA:DNA hybrids *in vitro*, the prevailing model posits that SETX uses its helicase activity to directly resolve R-loops. However, it remains possible that SETX rather controls R-loop formation by acting on RNAPII genomic occupancy. Indeed, SETX has been implicated in RNAPII transcription termination (38, 60–62) although it is unclear whether SETX has retained the ability of its yeast orthologue to induce termination or whether it rather plays an accessory role in this process.

The biological relevance of SETX is further underscored by the link between SETX mutations and two neurodegenerative diseases: ataxia with oculomotor apraxia (AOA2) and a juvenile form of amyotrophic lateral sclerosis dubbed ALS4 (63–69). While AOA2 is an autosomal recessive disease associated with SETX loss of function, ALS4 is an autosomal dominant disease provoked by toxic gain-of-function mutations in SETX (70). Several studies have reported decreased levels of R-loops in cells from ALS4 patients (25, 71–73), however it remains unclear whether ALS4 mutations alter the catalytic activity of SETX and how such alteration could eventually have an impact on R-loop levels.

Because to date there is no direct biochemical characterisation of SETX activities, much uncertainty exists about the molecular function of SETX. Here, we wished to elucidate the precise role of SETX in the control of R-loop accumulation and in transcription termination. We have purified the helicase domain of SETX and characterised its catalytic activities *in vitro*. We have included in our experiments the helicase domain of Sen1 to assess the conservation of the different activities, as well as potential divergences that may have appeared during evolution. We demonstrate that *in vitro*, SETX is an efficient nucleic acid-dependent ATPase and that it unwinds duplex substrates with 5′-3′ polarity, similarly to Sen1. Interestingly, we have also uncovered several differences between SETX and Sen1. Notably, SETX, unlike Sen1, hydrolyses ATP more efficiently in the presence of RNA than DNA, likely due to a more dynamic nature of the interaction of SETX with the RNA compared to the DNA. Moreover, SETX, but not Sen1, seems to have evolved a clear preference for R-loops over other types of nucleic acid substrates. Importantly, we show that SETX has retained the ability of Sen1 to induce transcription termination and that this function requires specific properties or features of mammalian RNAPII.

Finally, we have taken advantage of our biochemical tools to assess the effect of disease-associated mutations on SETX properties and catalytic activities. We have found that the AOA2-associated substitution (L1976R), when introduced in the helicase domain of SETX, impinges upon protein folding, which is in line with the SETX loss-of-function phenotype observed in this disease. In addition, we have found that the R2136H substitution, linked to ALS4, reduces the thermostability of the SETX helicase domain, with no discernible effect on its biochemical activities.

Taken together, our findings represent a major progress in the understanding of the function of SETX in the control of R-loop abundance and transcription termination and provide a framework to approach the molecular origin of neurodegenerative diseases linked to SETX mutations.

## MATERIAL AND METHODS

### Artificial intelligence-based modelling and data visualisation

AlphaFold 2.2 was used to generate a three-dimensional model of the helicase domain of SETX (amino acids 1699-2454) (74). Molecular graphic images were produced using the UCSF ChimeraX (75) package from the Resource for Biocomputing, Visualization, and Informatics at the University of California, San Francisco (supported by NIH P41 RR-01081).

### Plasmid construction

A fragment of DNA containing the coding sequence of the putative helicase domain of human SETX (amino acids 1674-2677; SETX-HD), codon-optimised for expression in insect cells, was first cloned into plasmid 2BcT (pET His6 LIC cloning vector; addgene #37236) to generate a C-terminally tagged SETX-HD-(His)_6_. Subsequently, SETX-HD-(His)_6_ was subcloned into plasmid 438A (pFastBac cloning vector with BioBrick PolyPromoter LIC Subcloning, addgene #55218) for purification of SETX-HD-(His)_6_ from insect cells. Plasmids 2BcT and 438A were purchased directly from QB3 Macrolab (UC Berkeley). Point mutations resulting in the following amino acid substitutions: K1969R; L1976R; DE2181, 2182AA; R2136H were introduced by site-directed mutagenesis. Oligonucleotides used are listed in Supplementary Table S1. The sequence integrity of all constructs was verified by sequencing.

### Protein expression in insect cells

To generate viruses enabling the production of proteins in insect cells, the coding sequences and the necessary regulatory sequences of the aforementioned constructs were transposed into bacmids using the *E. coli* strain DH10bac. Viral particles were obtained by transfection of the corresponding bacmids into the Sf9 cells using the FuGENE Transfection Reagent and further amplification. Proteins were expressed in 600 ml of Hi5 cells (infected at 1×10^6^ cells/ml) with the corresponding P1 virus at multiplicity of infection (MOI) >1. Cells were harvested 48 hours post-infection, washed with 1x PBS, and stored at −80 °C.

### Protein purification

Thawed cells were resuspended in lysis buffer (50 mM Tris-HCl, pH 8; 0.5 M NaCl; 10 % (v/v) glycerol; 1 mM DTT; 0.4 % (v/v) Triton-X; 10 mM imidazole) containing protease inhibitors (0.66 μg/ml pepstatin, 5 μg/ml benzamidine, 4.75 μg/ml leupeptin, 2 μg/ml aprotinin) and 25 U benzonase per ml of lysate. The cleared lysate was passed through 2 mL of Ni-NTA beads (Qiagen), equilibrated with 50 mM Tris-HCl, pH 8; 500 mM NaCl; 10 mM imidazole; and 1 mM DTT. SETX-HD was eluted with an elution buffer containing 50 mM Tris-HCl, pH 8; 350 mM NaCl; 1 mM DTT and 400 mM imidazole. The elution fractions containing SETX-HD were diluted 2-fold with 50 mM Tris-HCl, pH 8; 1 mM DTT and loaded onto a 1 mL HiTrap Heparin HP column (Cytiva). Proteins were eluted with 50 mM Tris-HCl, pH 7.5; 1 mM DDT and a 30-column-volume (CV) gradient of 200 to 650 mM NaCl. Fractions containing homogeneous SETX-HD were concentrated, and glycerol was added to a final concentration of 10 % before they were snap-frozen in liquid nitrogen and stored at −80 °C.

Recombinant Sen1-HD was expressed from the T7 promoter in *E. coli* and purified by consecutive Ni-affinity chromatography, heparin-affinity chromatography and gel filtration as described in (50).

The ten-subunit RNAPII core enzyme was purified from *S. cerevisiae* strain BJ5464 (76) by Ni-affinity chromatography followed by anion exchange on a Mono Q column essentially as previously described (77). Recombinant (His)_6_-tagged Rpb4/7 heterodimer was purified from *E. coli* by Ni-affinity chromatography followed by gel filtration as described in (77).

The mammalian, twelve-subunit RNAPII complex was purified from the calf (*Bos taurus*) thymus. Briefly, chromatin-bound RNAPII was purified to homogeneity by successive rounds of ion-exchange, immuno-affinity, ion-exchange, and size-exclusion chromatography as described in (78).

### Synthetic nucleic acid substrates

Oligonucleotides for preparing synthetic fluorescently labelled (Cy3 or FITC) RNA/DNA substrates were purchased from Sigma (HPLC purified) and their sequences are available in Supplementary Table 1. Duplex substrates were mixed in a 1:2 ratio of labelled versus unlabelled oligonucleotide and triplex substrates were prepared by mixing labelled molecules with a 5-fold excess of the unlabelled oligonucleotides in the annealing buffer (25 mM Tris-HCl, pH 7.5, 100 mM NaCl, 3 mM MgCl_2_), followed by initial denaturation at 75 °C for 5 min and subsequent cooling to room temperature. The annealed substrates were purified from a polyacrylamide native gel. Schematic representation of the substrates used is shown in Supplementary Figure S1.

### ATPase assays

The ATPase activity of SETX-HD and Sen1-HD proteins was assessed at 37 °C in 10 μl reactions containing 10 mM Tris–HCl, pH 7.5, 75 mM NaCl, 1 mM MgCl_2_, 1 μM ZnCl_2_, 1 mM DTT, 10 % (v/v) glycerol, 50 ng/μl of single-stranded DNA (ssDNA; oligo DL3701) or single-stranded RNA (ssRNA; oligo DL3316), 100 μM ATP-Mg, and 0.25 μM of 3000 Ci/mmol α^32^P-ATP. After preincubation of the mixtures for 5 min at 37 °C, the reactions were started with the addition of SETX-HD-WT (10 nM as the final concentration for reactions in the presence of ssDNA and 2.5 nM for reactions containing ssRNA), SETX-HD-DE/AA (25 nM as the final concentration) or Sen1-HD proteins (2 nM as the final concentration). Aliquots were taken at indicated time points, mixed with 2.5 volumes of quench buffer containing 10 mM EDTA and 0.5 % (w/v) SDS, and then subjected to thin-layer chromatography on PEI cellulose plates (Merck) in 0.35 M potassium phosphate pH 7.5. Plates were exposed to phosphorimager screens and scanned with a Typhoon scanner (GE Healthcare). The different radioactive species were analysed using the ImageQuant TL software (GE Healthcare).

### Electrophoretic Mobility Shift Assay (EMSA)

The indicated amounts of the different versions of SETX-HD and Sen1-HD proteins were incubated with fluorescently labelled nucleic acid substrates (at a final concentration of 10 nM) in buffer D (25 mM Tris-HCl, pH 7.5, 1 mM DTT, 0.1 mg/ml BSA, 5 mM MgCl_2_ and 100 mM NaCl and for nucleic acid substrates containing RNA, 2 U/μl of RNasin Ribonuclease Inhibitor (Promega)) for 20 min on ice. Loading buffer (60 % (v/v) glycerol in Orange-G) was added to the reaction mixtures and the samples were loaded onto a 7.5 % (w/v) polyacrylamide native gel in 0.5 x TBE buffer and run at 75 V for 1 hour. The different nucleic acid substrates were visualised using a FLA-9000 Starion scanner and quantified with the MultiGauge software (Fujifilm). To calculate the relative amount of bound nucleic acid substrate the band intensity (using the *band intensity - background* option) of the control sample, without protein, was set as 0 % of bound protein and the percent of bound protein for individual concentrations was calculated according to the control. Nucleic acid binding affinity graphs were generated with Prism-GraphPad 9 using the non-linear regression fit function.

### Helicase assays

The indicated amounts of the different versions of SETX-HD and Sen1-HD were incubated with fluorescently labelled nucleic acid substrates (at a final concentration of 10 nM) in buffer H (25 mM Tris-HCl, pH 7.5, 1 mM DTT, 50 mM NaCl, 20 mM creatine phosphate, 20 μg/ml creatine kinase, 3.5 mM MgCl_2_, and 2 mM ATP (for nucleic acid substrates containing RNA, 2 U/μl of RNasin Ribonuclease Inhibitor (Promega)) for 30 min at 37 °C. In reactions with duplex substrates a 5-molar excess of competitor oligonucleotide (pRMS680 for blunt-end, pRMS681 for 3′-overhang and pRMS682 for 5′-overhang RNA:DNA hybrids) was added to trap the unwound, unlabelled oligonucleotide. The reactions were stopped by adding a mixture of 2.5 % (w/v) SDS and 0.5 mg/ml proteinase K and incubated for 3 min at 37 °C. Loading buffer was added, and samples were loaded onto a 10-12 % (w/v) polyacrylamide native gel in 1x TBE and run at 80 V for 1 hour. Fluorescent nucleic acid species were visualised using a FLA-9000 Starion scanner and quantified by MultiGauge software (Fujifilm). Nucleic acid unwinding curves were generated by plotting the *percentage of unwound product* in Prism-GraphPad 9.

### Biolayer-interferometry (BLI)

To determine the kinetics of SETX-HD and Sen1-HD association and dissociation to ssDNA and ssRNA, respectively, BLI experiments were performed using a single–channel BLItz instrument in Advanced Kinetics mode (ForteBio) at room temperature, with shaking at 2200 rpm. Prior to the measurements, the streptavidin biosensors (SAX, ForteBio) were pre–hydrated in BLI buffer (buffer D, supplemented with 0.05 % (v/v) Tween 20) for 10 min. Biotinylated ssDNA (pRMS739) and ssRNA (pRMS740), respectively, were immobilised on the biosensor tip at a final concentration of 12 nM (oligonucleotide sequences can be found in Supplementary Table 1). The RNA substrate was diluted in BLI buffer supplemented with a RNase inhibitor at a final concentration of 2 U/μl (RNasin Ribonuclease inhibitor (Promega)). The output data are shown in a sensorgram as a change in optical thickness (in nm) induced by the binding of protein to the immobilised nucleic acid substrate over time. The protein association was measured at three different concentrations (12.5, 25, and 50 nM) for 180 s. Protein dissociation from the nucleic acid substrate was measured for 120 s by subsequent washing of the biosensor with BLI buffer. The association and dissociation constants were calculated in Prism GraphPad 9 software using the “Association then dissociation” function of non-linear regression in global analysis mode (constraint HotNM – shared value for all datasets) from triplicate measurements of each concentration for each protein with a coefficient of determination (R^2^) indicating the appropriateness of the fit.

### *In vitro* transcription termination (IVTT) assays

Termination assays were performed as previously described (77) with some modifications. Briefly, ternary elongation complexes (ECs) were assembled in a promoter-independent manner by first annealing a fluorescently labelled RNA (oligonucleotide DL2492) with the template DNA (oligonucleotide DL3352) and subsequently incubating the RNA:DNA hybrid (typically 1 pmol per reaction) with purified RNAPII (typically 250 ng per reaction) for 10 min at 20 °C. Next, a 2-fold excess of non-template (NT) strand (oligonucleotide DL3353) was added to the mixture and incubated for an extra 10 min at 20 °C. For reactions with yeast RNAPII we also added 2 pmol of recombinant Rpb4/7 and incubated the mixtures for additional 10 min. Ternary ECs were then immobilised on streptavidin beads (Dynabeads MyOne Streptavidin T1 from Invitrogen) and washed with transcription buffer (TB, 20 mM Tris–HCl, pH 7.5, 100 mM NaCl, 8 mM MgCl_2_, 10 μM ZnCl_2_, 10 % (v/v) glycerol, and 2 mM DTT) containing 0.1 % (v/v) Triton-X, TB containing 0.5 M NaCl, and finally TB. IVTT reactions were performed in TB in a final volume of 20 μl in the absence or the presence of 100 nM Sen1-HD or 600 nM SETX-HD. Transcription was initiated after addition of a mixture of ATP, UTP, and CTP (1 mM each as the final concentration in the reaction) and incubated for 10 min at 37 °C. Reactions were then stopped by the addition of 1 μl of 0.5 M EDTA and the beads and supernatant fractions were separated. Beads fractions were directly resuspended in 8 μl of loading buffer (1× Tris-borate-EDTA, 8 M urea) whereas transcripts in the supernatant fractions were ethanol precipitated and resuspended in 8 μl of loading buffer. Samples were boiled for 5 min at 95 °C and subjected to 10 % (w/v) denaturing PAGE containing 8 M urea. Gels were scanned with a Typhoon scanner (GE Healthcare) and analysed with the ImageQuant TL software (GE Healthcare).

### Differential scanning fluorimetry

The thermal stability of SETX-HD WT and the R2136H variant was measured using the Prometheus NT.38 instrument (NanoTemper Technologies). NanoDSF grade capillaries were filled with protein samples at 1 mg/ml in protein dilution buffer (25 mM Tris-HCl, pH 7.5, 150 mM NaCl, 0.01 % (v/v) NP-40, and 1 mM DTT). Protein unfolding was detected at a temperature range of 20 to 95 °C with a 1 °C/min heating rate and 90 % excitation power. Transition points (onset of protein unfolding (T_o_) and protein melting temperature (T_m_)) were determined from the ratio of tryptophan emission at 330 and 350 nm. The temperature of protein aggregation onset (T_agg_) was determined by light scattering, which is the back-reflected light intensity passing through the sample.

## RESULTS

### Expression and purification of the putative helicase domain of SETX

To investigate the mechanism of action of SETX we adopted an *in vitro* approach consisting in the purification and characterisation of the putative helicase domain of SETX (henceforth named SETX-HD), located at the C-terminal region of the protein comprising amino acids (aa) 1699-2454 (Figure 1A). AlfaFold2 predicts that SETX-HD adopts a fold similar to its yeast orthologue Sen1, with a notable difference in the length of the subdomain 1C (the ‘prong’) (Figure 1B), suggesting that it may confer different properties to SETX-HD (see discussion).

**Figure 1.**
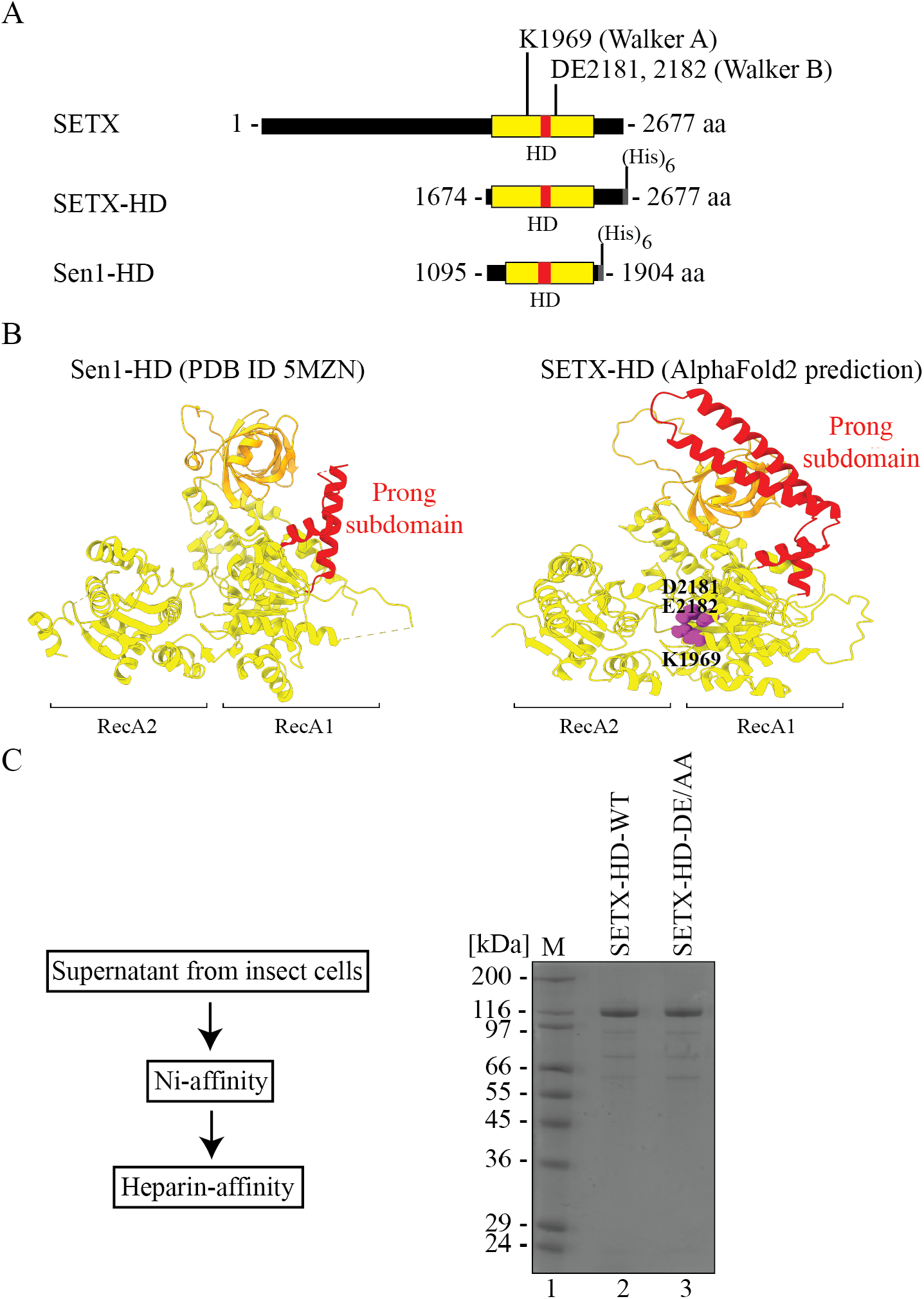
Design and purification of the helicase domain of senataxin. **(A)** Design of a variant of senataxin (SETX; amino acids (aa) 1674-2677), containing the putative helicase domain (HD, depicted in yellow), located between aa 1699 and 2454. A subdomain of the helicase domain, called the “prong” is depicted in red. A (His)_6_-tag is fused to the C-terminus of the fragment. The positions of Walker A and Walker B motifs are indicated. A fragment of Sen1 (Sen1-HD, aa 1095-1904), containing its helicase domain, used as a reference, is depicted below. **(B)** Comparison of the structure of Sen1 helicase domain (PBD ID 5MZN -(50)) and the helicase domain of SETX, predicted by AI-mediated modelling using AlphaFold2. RecA1 and RecA2 domains are depicted in yellow, β-barrel in orange, and the “prong” subdomain in red. In purple are highlighted the key residues responsible for ATP hydrolysis (K1969, within the Walker A motif), as well as for Mg^2+^ coordination (D2181, E2182, within the Walker B motif). **(C)** Procedure and representative results of purification of SETX-HD. Left: Cleared supernatant from insect cells was applied onto a Ni-affinity chromatography column and subsequently onto a heparin-affinity chromatography column. Right: Coomassie-stained SDS-PAGE gel showing purified SETX-HD variants (WT: wild-type and DE/AA: DE2181,2182AA). M: molecular weight marker.

The N-terminal boundary of the investigated variant was set at aa 1674, thereby including amino acids corresponding to a key structural feature of the helicase domain of Sen1 (the yeast orthologue of SETX; henceforth named Sen1-HD) termed the “brace”, which fastens three additional structural features within the helicase core: RecA1, the “stalk”, and the “barrel” and is likely critical for proper protein folding (50). A hexa-histidine tag was placed on the C-terminus of SETX-HD to avoid possible interference with the folding of this domain, as well as to enrich the full-length protein during the purification. To gain an evolutionary perspective of the activities of SETX-HD, we included in our experiments Sen1-HD (aa 1095-1904), which has been extensively characterised previously (Supplementary Figure S2A (49, 50)).

Two SETX-HD variants, mutated either in the Walker A motif, responsible for ATP hydrolysis (K1969R), or in the Walker B motif, responsible for Mg^2+^ binding (DE2181, 2182AA), were generated to determine the role of ATP hydrolysis in the potential activity(ies) of SETX-HD.

The wild-type (WT) version of SETX-HD (SETX-HD-WT), as well as the mutants in Walker A and Walker B motif, respectively, were expressed in insect cells, but only SETX-HD-WT and the mutant in the Walker B motif (hereafter SETX-HD-DE/AA) were soluble enough to be purified to near homogeneity (Figure 1C; Supplementary Figure S2B), which was necessary for subsequent analyses.

### The helicase domain of SETX exhibits a preference for RNA over DNA

Initially, we aimed to characterise the ATPase activity of purified SETX-HD. As helicases strictly require the interaction with nucleic acids to hydrolyse ATP, we first investigated the binding of SETX-HD and Sen1-HD to single-stranded nucleic acid substrates using electrophoretic mobility-shift assay (EMSA). Sen1-HD bound both ssDNA and ssRNA with similar affinity, in agreement with the existing literature (Figure 2A, Supplementary Figure S3A) (48, 49). In contrast, SETX-HD-WT bound more readily ssDNA than ssRNA (at 12.5 nM SETX-HD bound 33.6 % of the ssDNA substrate, whilst 19.4 % of the ssRNA substrate; at 25 nM SETX-HD bound 71.5 % ssDNA, whilst 35.9 % ssRNA) (Figure 2A, Supplementary Figure S3A). The SETX-HD DE/AA variant had the same nucleic acid binding pattern as SETX-HD-WT (Supplementary Figure S3A, S3B).

**Figure 2:**
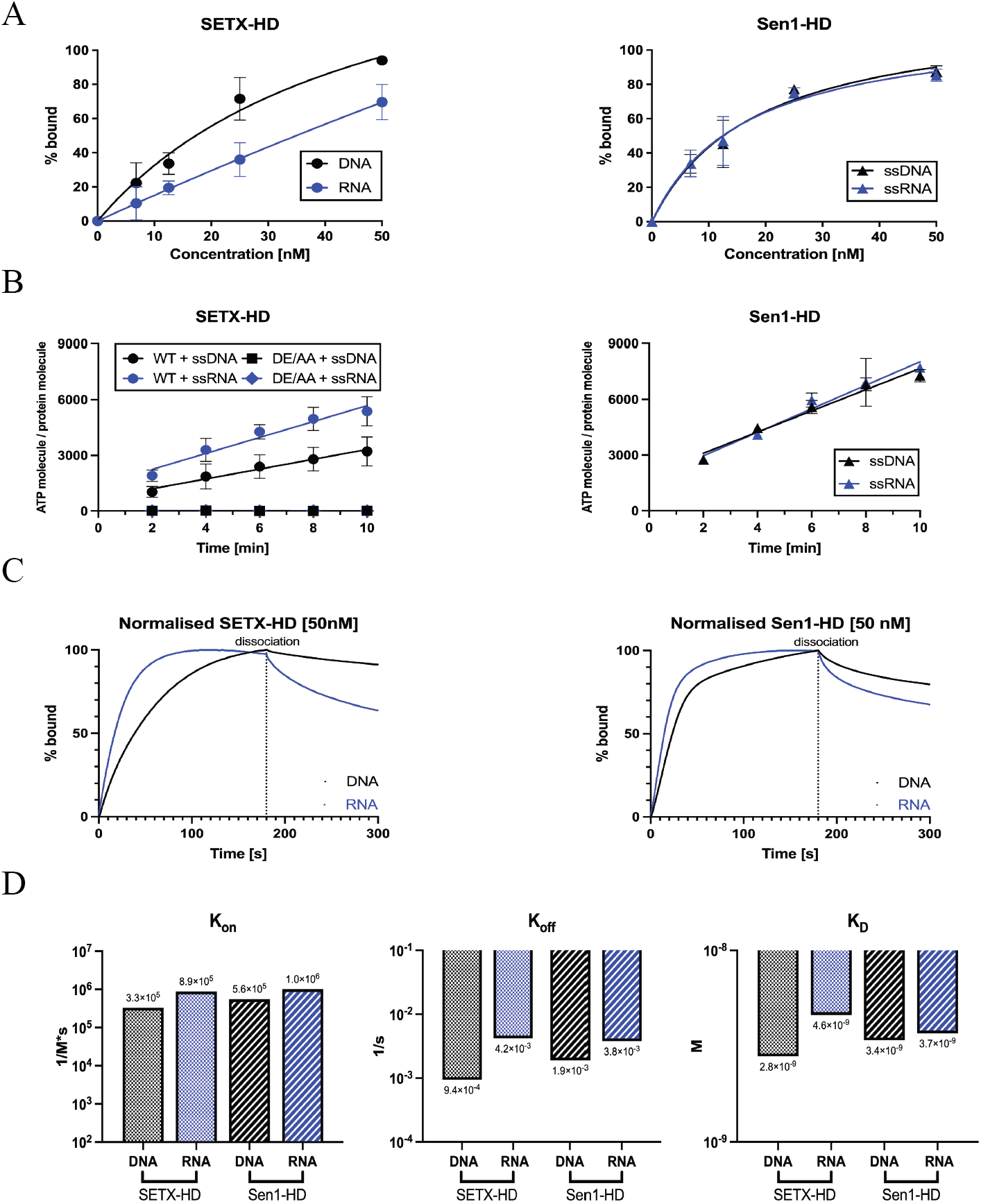
SETX, in contrast to Sen1, hydrolyses ATP more efficiently in the presence of single-stranded RNA than DNA. **(A)** Quantification of electrophoretic mobility-shift assays (EMSA) with SETX-HD and Sen1-HD, respectively, and single-stranded DNA (pRMS679, in black) or single-stranded RNA (pRMS678, in blue). The sequences of the different nucleic acid substrates are indicated in Supplementary table S1. The data points in each graph represent the mean of 3 independent experiments (except for SETX-HD + ssDNA: n = 4), error bars represent the standard deviation (SD). A scan of a gel of a representative experiment is shown in Supplementary Figure S3A. **(B)** Quantification of the amount of ATP hydrolysed by SETX-HD-WT, SETX-HD-DE/AA, and Sen1-HD, respectively, in the presence of single-stranded DNA (DL3701) or RNA (DL3316), respectively, plotted as a function of time. The data points represent the mean of 3 independent experiments, error bars represent the standard deviation (SD). A scan of a representative experiment is included in Supplementary Figure S3C. **(C)** SETX-HD dissociates from RNA more rapidly than from DNA. Sensorgrams from bio-layer interferometry (BLI) assay representing the mean of 3 measurements for each condition. Twelve fmoles of biotinylated ssDNA (pRMS739) and ssRNA (pRMS740), respectively, were bound to a SAX biosensor. SETX-HD (50 nM) and Sen1-HD (50 nM), respectively, were then submerged into the biosensor, to measure protein association for 180 s. Protein dissociation from nucleic acids was measured in BLI buffer for 120 s. Sensorgrams depict the comparison between association and dissociation rates of SETX-HD (left) and Sen1-HD (right), respectively, to DNA and RNA. Binding curves were normalised in Prism GraphPad 9. **(D)** Quantification of bio-layer interferometry (BLI) assays measuring the rate of association and dissociation of SETX-HD and Sen1-HD, respectively, to immobilised biotinylated single-stranded DNA and RNA. Graphs represent the association (K_on_), dissociation (K_off_), and dissociation (K_D_) constants, respectively, for SETX-HD and Sen1-HD with DNA (in black) and RNA (in blue). The data points represent the values calculated in Prism GraphPad 9 using the “Association then dissociation” function under global analysis mode from triplicate measures of three concentrations (12.5 nM, 25 nM, and 50 nM) for each condition. Sensorgrams for individual concentrations can be found in Supplementary Figure S3D.

In agreement with the EMSA assays and the existing literature (48), both ssDNA and ssRNA stimulated ATP hydrolysis by Sen1-HD to a similar extent (Figure 2B; Supplementary Figure S3C). Intriguingly, ssRNA stimulated the ATPase activity of SETX-HD nearly 1.7-fold more efficiently than ssDNA. As expected, SETX-HD-DE/AA did not hydrolyse ATP in the presence of neither ssDNA nor ssRNA, indicating that the observed ATPase activity is inherent to SETX-HD and not to any potential contaminating activity (Figure 2B, Supplementary Figure S3C).

To gain mechanistic insight into the observed different behaviour of SETX-HD and Sen1-HD, we employed biolayer-interferometry (BLI), which assesses the interaction between proteins and nucleic acids in real-time, enabling us to determine the kinetic parameters of the binding of SETX-HD and Sen1-HD to different nucleic acid substrates. We analysed the binding of SETX-HD and Sen1-HD to ssDNA and ssRNA immobilised on a biosensor (Figure 2C, Supplementary Figure S3D). SETX-HD associated with ssRNA 3-times faster than with ssDNA, whilst Sen1-HD associated only slightly faster with ssRNA than ssDNA (SETX: K_on_^RNA^=8.9×10^5^ M^-1^s^-1^; K_on_^DNA^=3.3×10^5^ M^-1^s^-1^; Sen1: K_on_^RNA^=1×10^6^ M^-1^s^-1^; K_on_^DNA^=5.6×10^5^ M^-1^s^-1^). SETX-HD dissociated 4-times faster from ssRNA than ssDNA (K_off_^RNA^=4.2×10^-3^ s^-1^; K_off_^DNA^=9.4×10^-4^ s^-1^), whilst Sen1-HD dissociated twice faster from ssRNA than from ssDNA (K_off_^RNA^=3.8×10^-3^ s^-1^; K_off_^DNA^=1.9×10^-3^ s^-1^) (Figure 2D). Taken together, Sen1-HD has comparable affinity towards ssRNA and ssDNA (K_D_^RNA^=3.7×10^-9^ M; K_D_^DNA^=3.4×10^-9^ M). In contrast, SETX-HD exhibits 2-fold higher affinity towards ssDNA than ssRNA (K_D_^RNA^=4.6×10^-9^ M; K_D_^DNA^=2.8×10^-9^ M) (Figure 2D). These data are consistent with the EMSA experiments and provide a mechanistic understanding of the observed differences in the rates at which ssDNA and ssRNA stimulate the ATPase activity of SETX-HD and Sen1-HD, respectively.

Collectively our data suggest that SETX-HD is indeed a potent ATPase, which, over the course of evolution, has adapted to hydrolyse ATP more efficiently on ssRNA, which is most likely caused by its increased rate of dissociation from ssRNA.

### SETX unwinds nucleic acid duplexes with a 5′-to-3′ polarity

Once we have established that the helicase domain of SETX has robust nucleic acid-dependent ATPase activity, we aimed to determine whether and with what polarity SETX-HD unwinds duplex substrates. To this end, we used a series of RNA:DNA duplex substrates harbouring a 5′ ssRNA overhang, a 3′ ssRNA overhang, or no overhang (see Supplementary Figure S1), which enabled us to unambiguously determine the polarity of the translocation of SETX-HD on nucleic acids. First, we analysed the ability of SETX-HD-WT to bind such substrates and determined that SETX-HD displayed similar affinity for hybrids containing a 5′ ssRNA or a 3′ ssRNA overhang, whilst it did not bind a duplex, in agreement with the notion that SETX-HD only binds single-stranded nucleic acids (Supplementary Figure S4A, B). Subsequently, we performed a series of helicase assays with the same substrates, and we found that SETX-HD unwound only the substrate containing a 5′ ssRNA overhang, albeit less efficiently than Sen1-HD. This unwinding activity required the presence of ATP, as well as the ability of SETX-HD to hydrolyse ATP, since the SETX catalytic mutant DE/AA was virtually inactive (Figure 3A, B).

**Figure 3:**
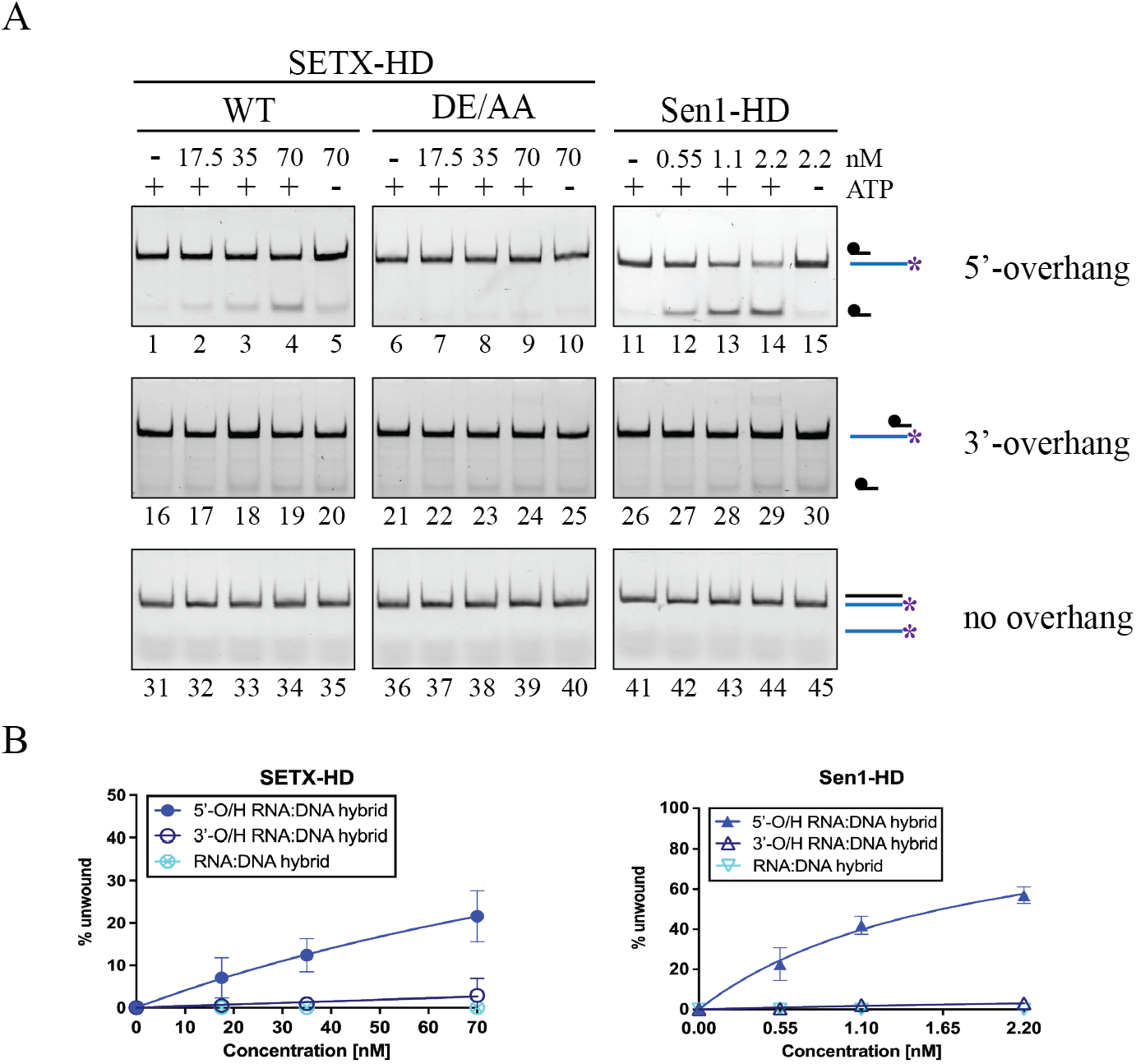
SETX, similarly to Sen1, can unwind RNA:DNA duplexes with a 5′-to-3′ polarity. **(A)** SETX-HD unwinds RNA:DNA duplexes with a 5′-overhang in an ATP dependent manner. Helicase assays were performed in the presence of the indicated amounts of SETX-HD-WT, SETX-HD-DE/AA, and Sen1-HD, respectively, and 10 nM fluorescently labelled RNA:DNA duplexes containing either a 5′ (lanes 1-15) or a 3′ (lanes 16-30) singlestranded overhang (O/H), or blunt ends (lanes 31-45). Where indicated, ATP was omitted. **(B)** Quantification of the experiments shown in (A). The data points in the graphs represent the mean of 3 independent experiments (for 5′-O/H RNA:DNA duplex, n = 4) and the error bars represent the standard deviation (SD).

Our data indicate that SETX-HD indeed translocates, in an ATP-dependent manner, on single-stranded nucleic acids with a 5′-to-3′ polarity.

### SETX-HD preferentially unwinds R-loop structures with a 5′ overhang

SETX has been implicated in the suppression of R-loop structures that form in cells during transcription (6, 38, 57–59). However, a direct, mechanistic evidence that SETX resolves R-loops is still missing. As SETX harbours a helicase domain, capable of unwinding RNA:DNA duplexes, we next decided to investigate whether SETX-HD can dismantle R-loop structures.

Initially, we performed EMSA experiments on R-loop structures with a 5′ ssRNA overhang (5′-O/H R-loop), a 3′ ssRNA overhang (3′-O/H R-loop), and with no overhang (R-loop) and found that SETX-HD bound the three types of molecules with similar affinity, probably because they all harbour at least one region of single-stranded nucleic acid (Supplementary Figure S5A, B). However, SETX-HD dismantled preferentially 5′-O/H R-loops in an ATP-dependent manner (Figure 4A-C; Supplementary Figure S5C, D), most likely by loading on the RNA and translocating from 5′ to 3′, in agreement with the results above (Figure 3). We observed a residual activity on the 3′-O/H R-loop substrate (Supplementary Figure S5C, compare lane 4 and lane 9), which is likely caused by partial, spontaneous opening of duplex regions allowing SETX-HD loading, rather than by a 3′-to-5′ translocation of SETX-HD on the substrate (see section above).

**Figure 4:**
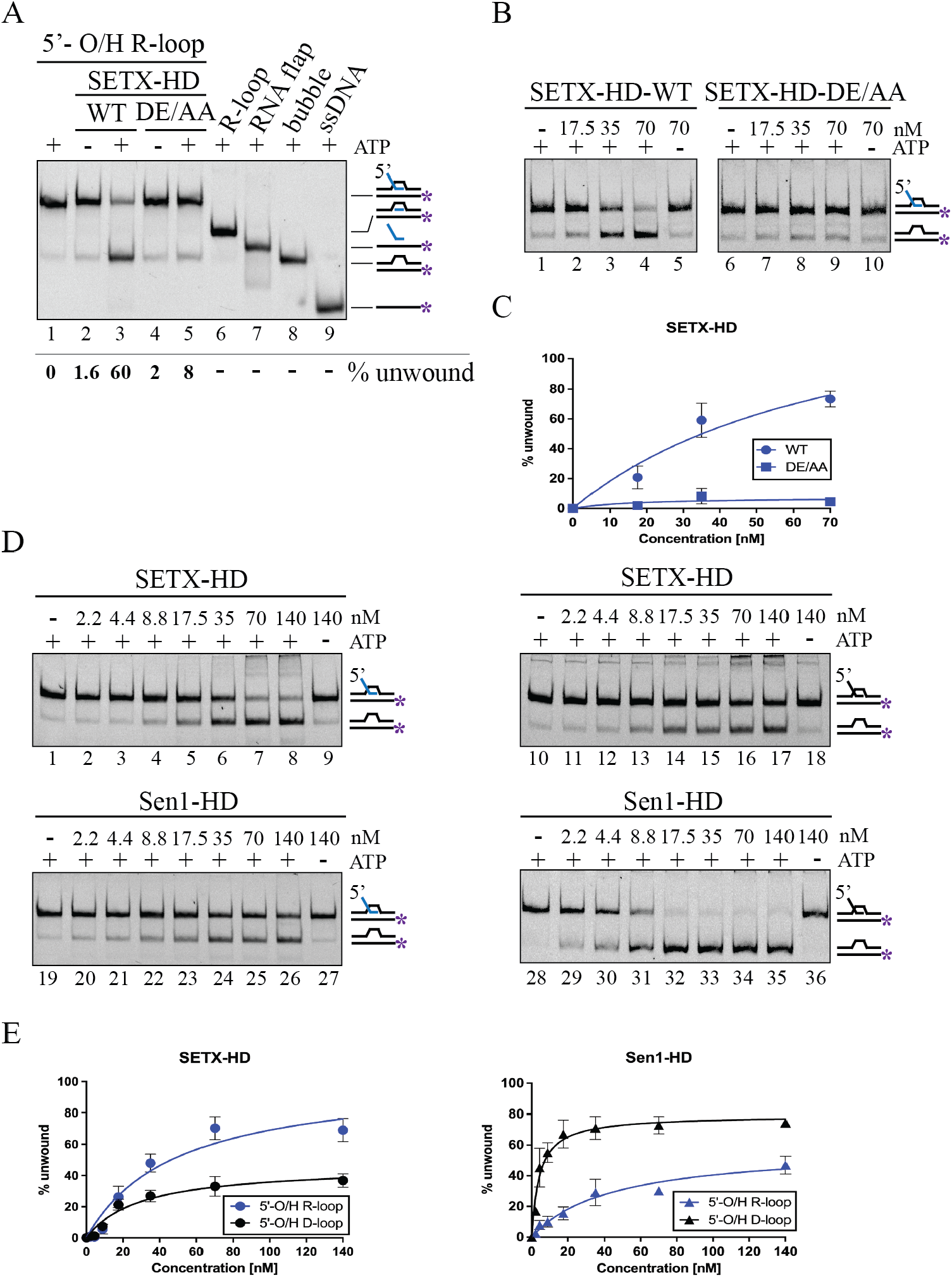
SETX-HD preferentially unwinds R-loop structures with a 5′-overhang. **(A)** SETX-HD unwinds the RNA:DNA duplex region of a 5′-O/H R-loop in an ATP-dependent manner. Helicase assays with 70 nM SETX-HD WT and DE/AA, respectively, and 10 nM fluorescently labelled 5′-O/H R-loop in the presence (lane 3 and 5) or absence (lane 2 and 4) of ATP. Lane 1 represents a control reaction with no protein. Samples containing all possible intermediates, resulting from either cleavage of ssRNA or SETX-mediated unwinding, were run on the gel for comparison. **(B)** SETX-HD unwinds 5′-O/H R-loops in a concentration-dependent manner. Increasing concentration of SETX-HD-WT and SETX-HD-DE/AA (17.5, 35, and 70 nM) were mixed with 10 nM fluorescently labelled 5′-O/H R-loop in the presence of ATP. Lanes 5 and 10 represent control experiments without ATP. **(C)** Quantification of the experiments shown in (B). The data points in the graph represent the mean of 5 (for WT + 5′-O/H R-loop) and 3 (for DE/AA + 5′-O/H R-loop) independent experiments and the error bars represent the SD. **(D)** SETX-HD preferentially unwinds 5′-O/H R-loop structures, whilst Sen1-HD preferentially unwinds 5′-O/H D-loop structures. Increasing concentrations of the indicated proteins were mixed with 10 nM fluorescently labelled 5′-O/H R-loop (lanes 1-9 SETX-HD; 19-27 Sen1-HD) or 5′-O/H D-loop (lanes 10-18 SETX-HD; 28-36 Sen1-HD). Where indicated, ATP was omitted. **(E)** Quantification of (D). Data points in graph represent the mean of 3 independent experiments and error bars represent the SD.

Next, we analysed the helicase activity of SETX-HD and Sen1-HD on both 5′-O/H D-loop and 5′-O/H R-loop structures. We found that SETX-HD unwound 5′-O/H R-loops significantly better than 5′-O/H D-loops (Figure 4D, E), although SETX-HD binds 5′-O/H R-loops and 5′-O/H D-loops with similar affinity (Supplementary Figure S5A, E, F). This result agrees with the observation that ssRNA stimulates the ATPase activity of SETX-HD more efficiently than ssDNA (Figure 2B; Supplementary Figure S3C). On the contrary, Sen1-HD unwound 5′-O/H D-loops more efficiently than 5′-O/H R-loops, which is in agreement with previous experiments conducted on RNA:DNA and DNA:DNA duplexes (48). Intriguingly, SETX-HD, unlike Sen1-HD, unwound triplex 5′-O/H R-loops 4-fold more efficiently than duplex 5′-O/H RNA:DNA hybrids (Supplementary Figure S5G, H), strengthening the notion that R-loops are the preferred substrate for SETX to dismantle.

In summary, our data suggest that SETX-HD, unlike Sen1-HD, dismantles R-loops more readily than D-loops if a 5′ overhang is present.

### SETX terminates transcription *in vitro* in a species-specific manner

In addition to the function in the resolution of R-loops, several studies assigned to SETX a role in transcription termination (38, 60, 79). It has been shown that *in vitro* Sen1 efficiently terminates transcription by translocating along the nascent RNA to dislodge RNAPII from the DNA (48, 51). Furthermore, Sen1-HD is sufficient to terminate transcription *in vitro* (48, 50). However, it remains unclear whether SETX has retained the capacity of its yeast orthologue to induce termination or whether it rather plays an indirect role in this process.

To address this question, we used an already established assay, initially developed to investigate the mechanism by which Sen1 terminates RNAPII-mediated transcription (48). In this assay, promoter-independent assembly of the elongation complex (EC) is performed using a 9 nt RNA:DNA hybrid, which occupies the catalytic centre of RNAPII (Figure 5A). Ternary ECs (yeast or mammalian) are attached to streptavidin beads via the 5′ biotin of the non-template strand, enabling subsequent separation of the bead-associated (B) and the supernatant (S) fractions. The RNA molecule is fluorescently labelled to allow its detection. The transcription template contains a G-less cassette followed by a G-stretch in the non-template strand. After adding a mixture of ATP, UTP, and CTP, RNAPII transcribes until it encounters the G-rich sequence. The ability of Sen1-HD and SETX-HD, to provoke dissociation of ECs paused at the G-rich stretch is then monitored by comparing the amount of nascent RNA released to the supernatant in the absence and the presence of these proteins in the reactions.

**Figure 5.**
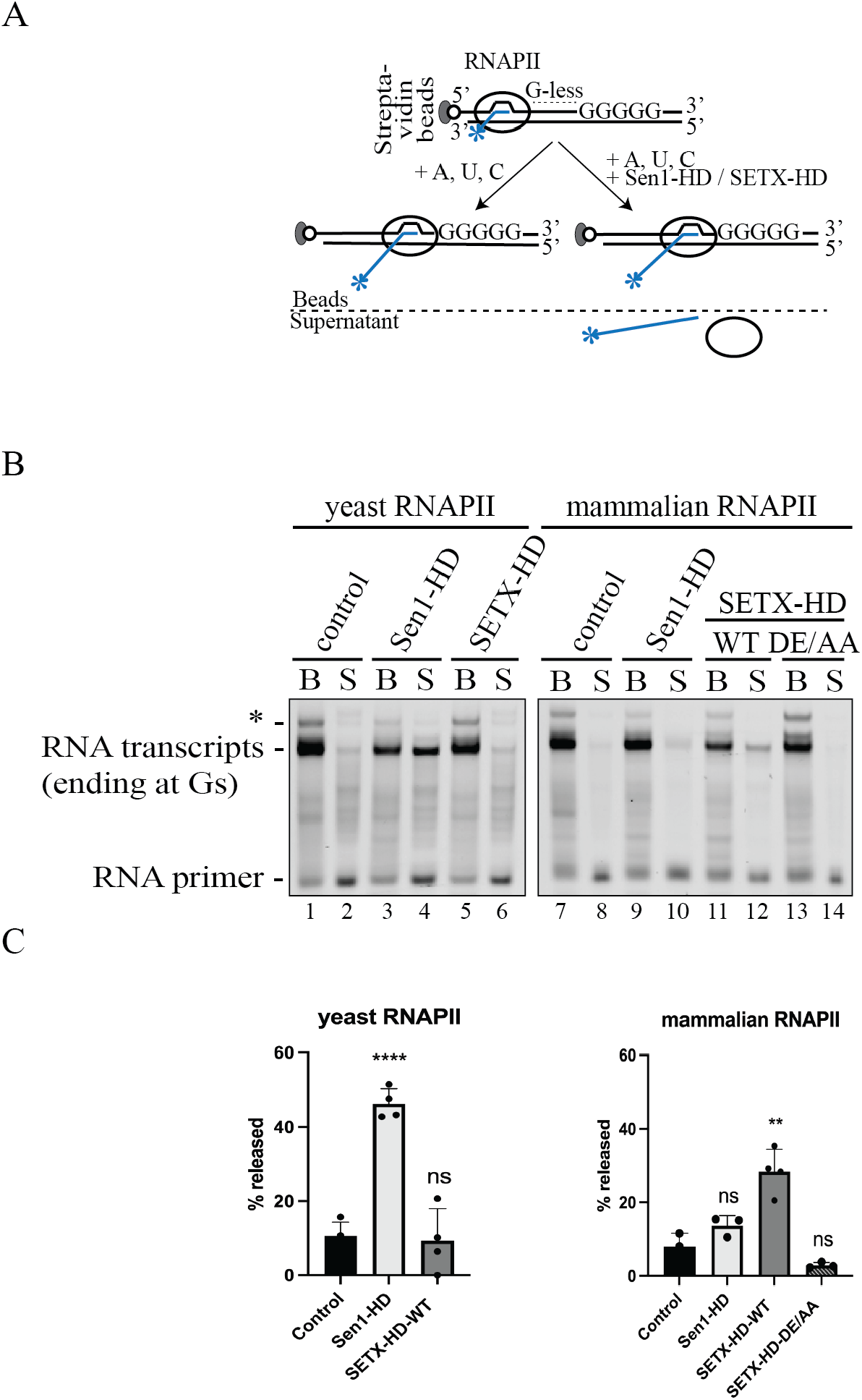
SETX-HD-WT terminates transcription *in vitro* in a species-specific manner. **(A)** Schematic representation of the *in vitro* transcription-termination (IVTT) assay (48). Promoter-independent assembly of the elongation complex (EC) is performed using a 9 nt RNA:DNA hybrid, which occupies the catalytic centre of RNAPII. Ternary ECs are attached to streptavidin beads via the 5′ biotin of the non-template strand, enabling subsequent separation of bead-associated (B) and supernatant (S) fractions. The RNA is fluorescently labelled with FAM at the 5′-end. The transcription template contains a G-less cassette followed by a G-stretch in the nontemplate strand. After adding an ATP, UTP, CTP mix, the RNAPII transcribes until it encounters the G-rich sequence. Subsequently, Sen1-HD/SETX-HD may provoke dissociation of ECs paused at the G-rich stretch and therefore the release of RNAPII and associated transcripts to the supernatant. **(B)** IVTT assay using either yeast RNAPII (lanes 1-6) or mammalian RNAPII (lanes 7-14). Transcript release from the bead fraction (B) to the supernatant fraction (S) was monitored upon addition of Sen1-HD, SETX-HD-WT, or SETX-HD-DE/AA, as indicated. Lanes 1-2 and 7-8 represent control reactions with no protein. RNA species were analysed by denaturing PAGE. The RNA transcripts associated with RNAPIIs paused at the G-stretch are indicated. Asterisk denotes transcripts associated with RNAPIIs that over-read the G-stretch due to small contamination of the nucleotide solutions with GTP. **(C)** Quantification of (B). Quantification of RNA transcript in the supernatant fraction as an estimate of the efficiency of termination. Values represent the mean of three independent experiments (for Sen1-HD with yeast RNAPII: n = 4) and error bars denote the SD. The values obtained for each protein were compared to the control using a two-tailed unpaired Student’s t-test. The resulting p-values (P) are represented by ns (nonsignificant) P > 0.05; ** P < 0.01, and **** P < 0.001.

First, we found that Sen1-HD efficiently dislodged yeast RNAPII from the DNA since ~50 % of the nascent RNA was present in the supernatant (Figure 5B, lane 4; Figure 5C), as previously shown (48). Under the same experimental set-up, however, SETX-HD did not promote the release of the polymerase and the nascent RNA from the DNA (Figure 5B, lane 6), indicating that either SETX has not retained the same ability of its yeast counterpart or that it displays a species-specific termination activity. To distinguish between these possibilities, we repeated the assay with mammalian RNAPII instead of yeast RNAPII. Interestingly, the addition of SETX-HD-WT, but not SETX-HD-DE/AA resulted in the release of the nascent RNA into the supernatant (Figure 5B, compare lane 12 and 14; Figure 5C), indicating that SETX may mediate the dissociation of mammalian ECs in an ATP-dependent manner. Strikingly, Sen1-HD could not promote the efficient release of the mammalian RNAPII and the associated RNA into the supernatant (Figure 5B, compare lane 4 and 10), indicating a specialisation for yeast RNAPII.

Collectively, these data support the notion that SETX-HD induces transcription termination, albeit less efficiently than Sen1-HD, and that this activity likely requires specific features of mammalian RNAPII.

### Biochemical characterisation of variants of SETX associated with neurodegenerative diseases

Our thorough biochemical characterisation of the SETX-HD enabled us to assess the effect of mutations within the helicase domain that are associated with neurodegenerative diseases. Two different diseases are associated with mutations in SETX: (i) Ataxia with oculomotor apraxia type 2 (AOA2); (ii) amyotrophic lateral sclerosis type 4 (ALS4). Whilst mutations causing AOA2 resemble SETX null phenotypes *in vivo* (63, 65–68, 72), mutations associated with ALS4 cause milder phenotypes (25, 64, 73, 80). We constructed and attempted to purify two mutant versions of SETX: SETX-HD-L1976R (SETX-HD-L/R), associated with AOA2 (67) and SETX-HD-R2136H (SETX-HD-R/H), associated with ALS4 (64) (Figure 6A). Leucine 1976 is located within the RecA1 subdomain, suggesting that the L1976R substitution may impinge upon the ATPase activity and/or the overall structural integrity of the domain, whereas arginine 2136 is located at the bottom of the ‘prong’ subdomain, suggesting a milder effect of the R2136H substitution on the overall activity and/or folding of the domain (Figure 6B).

**Figure 6.**
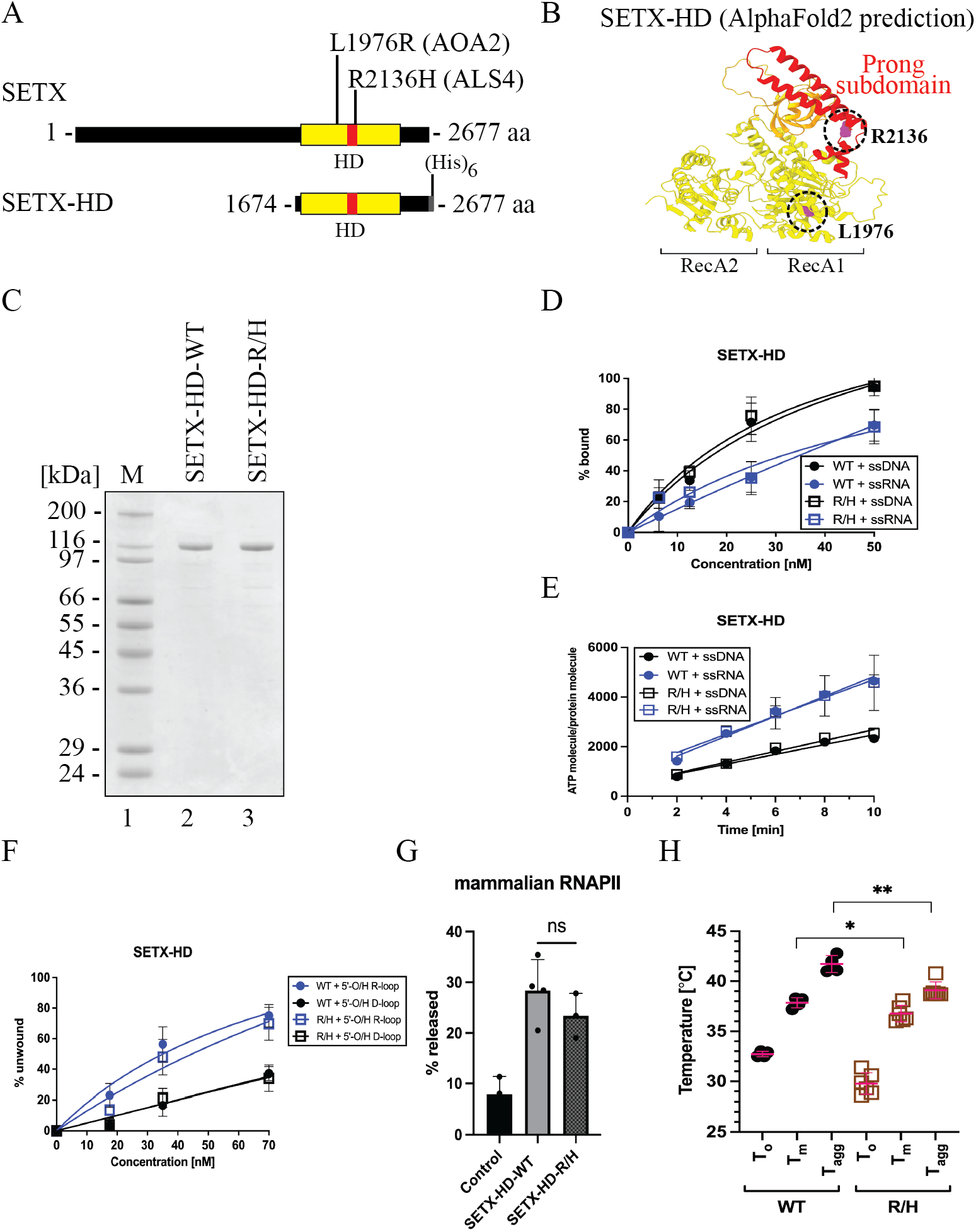
Purification and biochemical characterisation of SETX-HD variants associated with neurodegenerative diseases. **(A)** Schematic representation of SETX and position of the investigated mutations associated with AOA2 (the SETX-HD L1976R (L/R) variant) and ALS4 (the SETX-HD R2136H (R/H) variant). **(B)** Structure of the helicase domain of SETX predicted by AI-mediated modelling using AlphaFold2. RecA1 and RecA2 domains are depicted in yellow, β-barrel in orange, and the “prong” subdomain in red. In purple are highlighted the positions of the amino acids mutated in AOA2 (L1976) and ALS4 (R2136). **(C)** SDS-PAGE gel showing the purified proteins SETX-HD-WT and SETX-HD-R/H. M: molecular weight marker. **(D)** SETX-HD-R/H binds nucleic acid substrates to the same extent as SETX-HD-WT. Quantification of EMSA with SETX-HD-WT and SETX-HD-R/H, respectively, in the presence of single-stranded DNA (pRMS679, in black) or singlestranded RNA (pRMS678, in blue). The data points in the graph represent the mean of several independent experiments (WT + ssDNA: n = 4; WT + ssRNA: n = 3; R/H + ssDNA: n = 3 and R/H + ssRNA: n = 4), while error bars represent the SD. A scan of a representative experiment is included in Supplementary Figure S6C. **(E)** SETX-HD-R/H exhibits ATPase activity comparable to SETX-HD-WT. Quantification of ATP hydrolysis by SETX-HD-WT and SETX-HD-R/H in the presence of single-stranded DNA or RNA plotted as a function of time. The data points represent the mean of several independent experiments (WT/R/H + ssRNA: n = 4 and WT/R/H + ssDNA: n = 3), while error bars represent the SD. A scan of a representative experiment is shown in Supplementary Figure S6D. **(F)** SETX-HD-R/H unwinds 5′-O/H R-loop and 5′-O/H D-loop structures similarly to SETX-HD-WT. Quantification of helicase assays performed in the presence of increasing concentrations of SETX-HD-WT and SETX-HD-R/H with 5′-O/H R-loop (blue) or 5′-O/H D-loop (black) as substrates. The data points represent the mean of 4 (for 5′-O/H R-loop) or 3 (for 5′-O/H D-loop) independent experiments and error bars represent the SD. A scan of a representative experiment can be found in Supplementary Figure S6E. **(G)** SETX-HD-R/H is as efficient as SETX-HD-WT in terminating transcription *in vitro*. Quantification of RNA transcripts in the supernatant fraction. Values represent the mean of 3 or 4 (for SETX-HD-WT) independent experiments and error bars represent the SD. The values obtained for the mutant were compared to the WT using a two-tailed unpaired Student’s t-test. The resulting p-value (P) is indicated by ns (non-significant) P>0.05. A representative scan of the experiment can be found in Supplementary Figure S6F. **(H)** SETX-HD-R/H variant is less thermostable than SETX-HD-WT. Quantification of the thermostability of SETX-HD-WT and SETX-HD-R/H measured by nanoDSF. Plots represent the mean of replicate measures (WT: n = 4; R/H: n = 6) indicating the temperatures corresponding to the onset of protein unfolding (T_o_), the melting temperature (T_m_), and the onset of protein aggregation (T_Agg_). Error bars represent the SD. The p-value resulting from a two-tailed unpaired Student’s t-test is represented by * P < 0.05; ** P < 0.01. A representative thermograph can be found in Supplementary Figure S6G.

The SETX-HD-L/R variant could be expressed in insect cells, but it was insoluble (Supplementary Figure S6A), suggesting that the mutation affected the proper folding of the domain, thereby likely explaining the phenotype *in vivo*. On the other hand, SETX-HD-R/H was soluble and could be purified to near homogeneity (Figure 6C; Supplementary Figure S6B). We performed a complete set of biochemical experiments, comparing SETX-HD-R/H to the wild-type version. The behaviour of the SETX-HD-R/H variant was indistinguishable from that of the wild-type in terms of: (i) the binding to nucleic acid substrates (Figure 6D; Supplementary Figure S6C); (ii) ATP hydrolysis (Figure 6E; Supplementary Figure S6D); (iii) the helicase activity on both R-loop and D-loop substrates (Figure 6F; Supplementary Figure S6E); and (iv) the ability to induce transcription termination *in vitro* (Figure 6G; Supplementary Figure S6F). Finally, to monitor for possible differences in the thermal stability of SETX-HD-R/H, we conducted differential scanning fluorimetry. Indeed, we observed a statistically significant shift in the melting temperature (T_m_) and the onset of protein aggregation (T_Agg_) for SETX-HD-R/H (Figure 6H; Supplementary Figure S6G), suggesting that this mutant variant may be less stable *in vivo*.

Taken together, our biochemical characterisation of the catalytic domain of SETX has enabled us to unveil that SETX is a bona fide helicase and transcription termination factor and has provided tools to directly address the impact of disease-associated mutations on the properties and activities of this relevant protein.

## DISCUSSION

Over the recent years, the putative helicase SETX has emerged as a safeguard of gene expression and genome integrity. Studies conducted in a variety of cellular models, mainly based on diverse phenotypic characterisations upon SETX knock-down, have implicated SETX in critical processes such as transcription termination and the regulation of gene expression, the control of R-loop and DNA damage accumulation, and the regulation of transcription-replication conflicts. All functions attributed to SETX have been proposed to rely on its enzymatic properties as a helicase. However, despite the important role of SETX in several fundamental biological processes and the causal connection between SETX mutations and two devastating neurodegenerative diseases, the molecular function of SETX has remained elusive. In the present study, we fill this gap by providing the first purification and biochemical characterisation of the helicase domain of SETX, which is responsible for its catalytic activities.

Likely the most extensively studied aspect of SETX function is its involvement in R-loop homeostasis. As mentioned in the introduction, R-loops appear as a consequence of normal DNA metabolic activities, such as transcription, and participate in several physiological processes. Only when R-loops accumulate/persist, they become a threat for genome stability and transcription (21). One of the strategies to curb the accumulation of R-loops is their direct dismantling by helicases. Depletion of SETX most often results in an increase of R-loop levels (5, 6, 38, 57–60, 72, 80). This has led many authors to propose that SETX directly resolves R-loops by unwinding the RNA:DNA hybrid portion of R-loops, therefore promoting the restoration of the natural double-stranded DNA. However, transcription, which is the major source of R-loops, is also substantially altered by SETX depletion (60–62, 72). A couple of recent studies have shown that the perturbations of RNAPII genomic distribution provoked by SETX deletion/depletion are rather associated with R-loop losses at many loci (81, 82). Therefore, the former data did not allow discriminating between a direct role for SETX in the resolution of R-loops from an indirect role via the regulation of transcription.

Here, using purified SETX helicase domain and synthetic R-loops we provide direct evidence that SETX can efficiently dismantle R-loops if they possess a 5′ ssRNA overhang (Figure 4 and Supplementary Figure S5C, D). Complementary experiments indicate that such R-loop resolving activity involves the loading of SETX to the exposed single-stranded RNA and its translocation in the 5′ to 3′ direction to unwind the RNA:DNA hybrid region of the R-loop. This type of R-loop (*i.e*., having a 5′ ssRNA overhang) is likely the most frequently formed during transcription when the nascent RNA invades the DNA behind the transcribing RNAPII. Additionally, R-loops may be formed at the front of RNAPII upon polymerase backtracking, for instance in response to damaged bases in the template DNA (83). According to our biochemical analyses, this kind of R-loop, lacking a ssRNA portion, would not be a good substrate for SETX, since we detected virtually no activity on such structures (Supplementary Figure S5C, lanes 11-15). It is possible that R-loops resulting from backtracking are removed by RNAPII itself once it resumes elongation, as it has been observed *in vitro* (83). Alternatively, other R-loop resolving RNA:DNA helicases, such as BLM, DDX5, DDX21, that are active on R-loops with no overhangs *in vitro* (84–86) may release the path for the polymerase. We also observed a weak SETX unwinding activity on R-loops with a 3′ ssRNA overhang (Supplementary Figure S5C, lanes 6-10). However, we believe that this activity is most likely the result of transient, spontaneous opening of the duplex DNA, followed by SETX translocation in 5′-3′ direction along the DNA strand annealed to the RNA. Such opening might be favoured by the 3′ ssRNA overhang. Altogether, our data allow us to propose that R-loops forming behind the transcribing RNAPII are the major targets of SETX *in vivo* (Figure 7).

**Figure 7.**
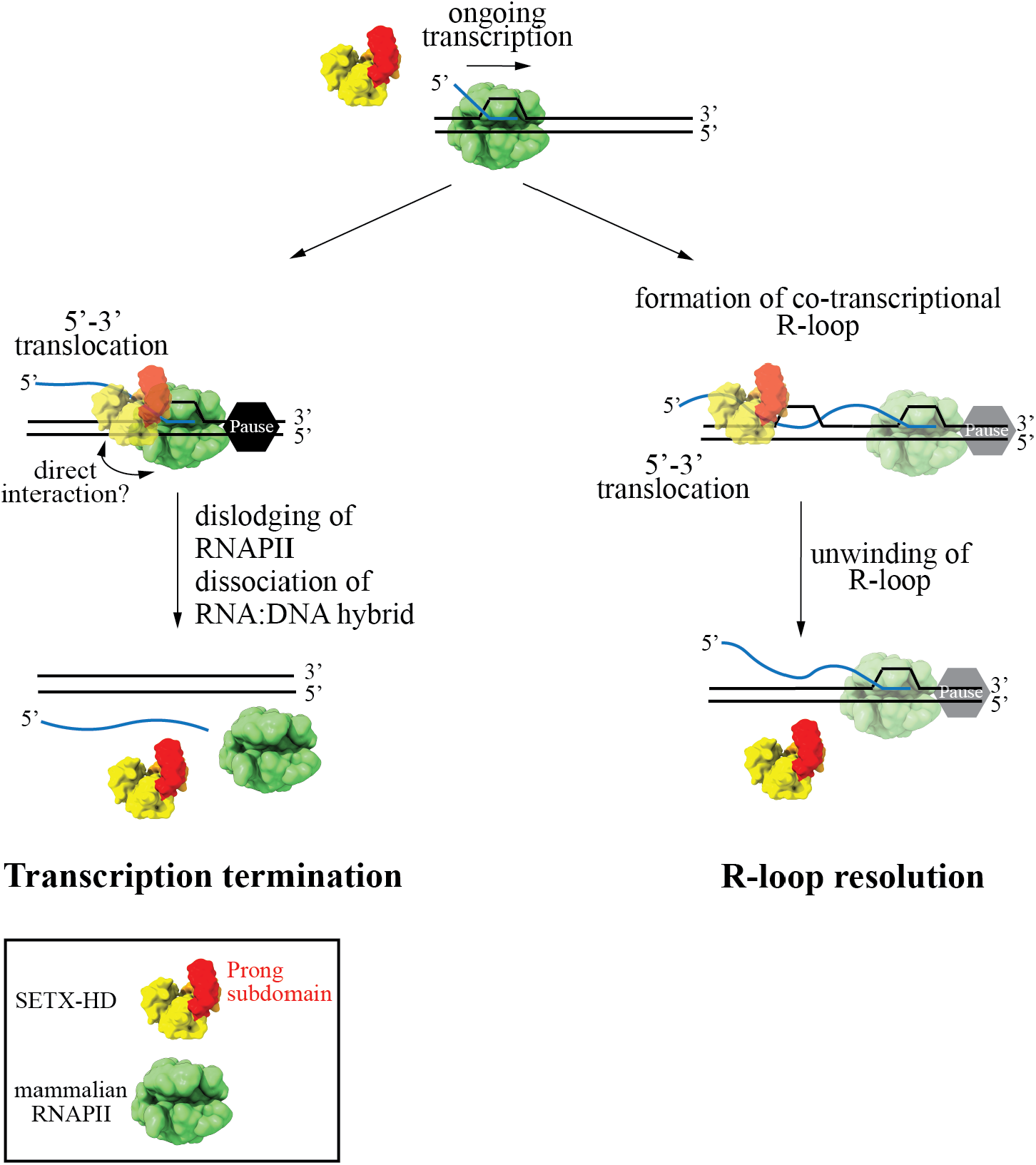
Schematic model of mechanisms by which senataxin may promote R-loop dissolution and transcription termination. RNAPIIs may pause/stall during transcription for instance upon reading specific DNA sequences or upon encountering other DNA-bound proteins or damaged template. SETX translocates on the single-stranded nascent RNA with a 5′-3′ polarity to dislodge RNAPII and its associated transcript. This process likely requires direct interactions between specific regions of SETX and RNAPII. Transcription termination by SETX may play a role in the regulation of gene expression and might also be important to prevent unscheduled persistence of RNAPIIs at TRC and DNA damage sites. R-loops may appear in various forms - with either a 5′, a 3′ RNA overhang, or possibly with no overhang. R-loops formed during transcription most often possess a 5′ RNA overhang. SETX can efficiently resolve this kind of structure by translocating in the 5′ −3′ direction on single-stranded RNA to disrupt the RNA:DNA hybrid within the R-loop and release the RNA moiety.

Initial functional studies of SETX uncovered its involvement in several facets of RNA metabolism including transcription termination (79). Later reports showed a role for SETX in the regulation of the expression of several genes involved in the establishment of circadian clock and in the antiviral immune response (61, 87). Specifically, SETX was proposed to promote premature transcription termination at those genes, in other words, to induce termination within the coding sequence leading to the production of truncated, non-functional mRNAs and, thus, gene downregulation. Additional studies also implicated SETX in transcription termination at the 3′ end of some genes (38, 60, 62). However, the function of SETX in transcription termination is often linked to R-loop metabolism since accumulation of both RNAPIIs and R-loops is observed at termination regions upon SETX depletion. Indeed, it has been proposed that SETX would not induce termination by itself but rather favour the action of XRN2, an exonuclease that plays a general role in transcription termination at protein-coding genes. By removing R-loops at termination regions, SETX would allow the access of XRN2 to the nascent transcript, since XRN2 is presumably unable to degrade the RNA hybridised to DNA; and thereby, XRN2 could ultimately reach the RNAPII and elicit its release from the DNA (38). Hence, the precise role of SETX in transcription termination has remained unclear.

In the present study we have used a highly purified *in vitro* system that has enabled us to assess the ability of SETX to induce transcription termination directly and separately from its activity on R-loops because the DNA templates, which we use do not form any R-loop (51). Importantly, we have demonstrated that, akin to its yeast homologue Sen1, SETX is a genuine transcription termination factor, which directly induces the release of paused RNAPIIs and associated transcripts from the DNA (Figures 5 and 7). We therefore propose that SETX uses its ability to dislodge RNAPII from the DNA to regulate gene expression and possibly to remove RNAPIIs from TRC sites, as recently showed for its yeast counterpart (56).

Interestingly, we have observed that SETX displays a clear transcription termination activity on bovine RNAPII, which is 99.9 % identical to human RNAPII (78) but not on the more divergent budding yeast RNAPII (Figure 5). Similarly, Sen1 works much more efficiently on its cognate RNAPII, indicating that these proteins have coevolved with RNAPII to remain transcription termination factors in their respective organisms. The species-specific character of SETX/Sen1 transcription termination activity may explain the inability of SETX to complement *sen1* deficiency in *S. cerevisiae* (88) and provides potentially relevant information about the precise mechanisms of termination. As outlined above, to promote termination, Sen1 and likely SETX translocates along the nascent transcript until it collides with RNAPII. However, it is unclear how such a collision results in the dislodging of RNAPII from the DNA. The fact that both proteins require species-specific features of RNAPII suggests that specific interactions between Sen1/SETX and RNAPII might play an important role in termination, adding an extra layer of complexity to this process. We speculate that such protein-protein interactions might induce destabilising conformational changes in the transcription elongation complex. We hypothesise that one of the involved interaction surfaces could be located at the prong because: (i) this subdomain plays an essential role in Sen1-mediated transcription termination both *in vitro* and *in vivo* (50); (ii) its location predicts that it would be in direct contact with the polymerase upon SETX-RNAPII collision; and (iii) remarkable differences can be appreciated between Sen1 and SETX prongs. Indeed, structure-guided sequence alignment shows that the prong domain of SETX is shorter than the one of Sen1 (86 aa vs. 104 aa) and there is poor sequence conservation (Supplementary Figure S7A). Whilst the central portion of the Sen1 prong (amino acids 1471–1538) is predicted to be disordered (50), the SETX structural model predicted by AplhaFold2 showed an additional large coiled-coil-like loop in the central region of the prong domain (Supplementary Figure S7B). Understanding, what are the structural determinants of SETX/Sen1 species-specific transcription termination activity is an interesting matter for future investigation.

Our side-by-side comparison of the biochemical properties of SETX and Sen1 has enabled us to look at the activities of these proteins from an evolutionary perspective. We have uncovered several notable differences between SETX and Sen1. Firstly, we have found that ssRNA stimulates the ATPase activity of SETX significantly more efficiently than ssDNA, which is not the case for Sen1 (Figure 2B and Supplementary Figure S3C). This is most likely due to a more dynamic interaction of SETX with ssRNA than with ssDNA (Figure 2C, D), which may favour a higher exchange of SETX on the RNA substrate and, therefore, support a higher ATP hydrolysis rate. We hypothesise that over the course of evolution, SETX has accumulated mutations on the RNA-binding surface conferring this specific behaviour. Using the available structure of the helicase domain of Upf1, a close homologue of Sen1/SETX, in complex with the RNA (PDB: 2XZO) (89) and taking advantage of the high conservation of the residues forming the helicase channel in the three proteins, we have modelled threading of the RNA substrate through Sen1 and SETX to rationalise their different nucleic acid binding properties (Supplementary Figure S7C). Interestingly, all residues contacting the sugar moieties are fully conserved between Sen1 and SETX, except for one at the entry channel, where the T1289 in Sen1 is replaced by V1884 in SETX (Supplementary Figure S7D, E). We hypothesise that the threonine in Sen1 may contact both DNA and RNA equally well, as the threonine residue can switch between polar and hydrophobic contacts to the ribose and deoxyribose, respectively. In contrast, SETX has a valine at this position, which disfavours interaction with the ribose moiety of RNA, thereby explaining the less stable binding of SETX to RNA compared to DNA (Figure 2 and Supplementary Figure S3A).

In addition, our data indicate that, unlike Sen1, SETX is particularly avid for R-loops since it unwinds triplex substrates (*i.e*., R-loops) more efficiently than duplex substrates (*i.e*., RNA:DNA hybrids) and is more active on R-loops than on D-loops (Figure 4D, E and Supplementary Figure S5G, H). It is therefore possible to envisage that SETX has evolved to play a more prominent role in R-loop resolution than Sen1.

Defects in transcription termination and R-loop metabolism have been implicated in the onset of several neurodegenerative diseases (25, 79, 80, 82, 90, 91), highlighting the importance of understanding these processes. Mutations in SETX are associated with ataxia with oculomotor apraxia (AOA2) and a juvenile form of amyotrophic lateral sclerosis dubbed ALS4 (63–69). AOA2 mutations lead to SETX loss of function and are associated with transcriptional stress, R-loop accumulation, increased oxidative stress, elevated levels of double-strand breaks and associated genome instability, similarly to the phenotypes observed upon SETX depletion (5, 69, 72, 79, 92–94). A previous study used Sen1 as a surrogate for SETX to assess the effect of a subset of AOA2-associated mutations on SETX biochemical activities and found that the mutations analysed are associated with improper protein folding, impaired RNA binding, or reduced ATPase activity (50). However, a number of SETX disease-associated mutations do not fall into positions conserved in the yeast homologue, notably those linked to ALS4, and the properties of Sen1 and SETX present some clear divergencies, which are important limitations of using Sen1 as a model to study SETX-associated diseases.

In the present study, we report a series of methods to purify the helicase domain of SETX and assess its relevant biochemical activities, which provides an ideal system to investigate the molecular effect of SETX disease-associated mutations located at this region. To validate these tools, we have characterised one AOA2 and one ALS4 mutation (Figure 6). Specifically, we have found that the L1976R substitution (67), causing AOA2, results in protein insolubility (Supplementary Figure S2B), which explains the loss-of-function phenotypes. In contrast, we have found that the ALS4-associated substitution R2136H (64) does not significantly affect protein solubility nor any of the measurable SETX activities (Figure 6). Expression of SETX mutant versions harbouring either the L389S or the R2136H ALS4 substitutions induce ALS-like phenotypes in mice models, supporting the idea that ALS4 is provoked by toxic SETX gain of function (70). The comparison of ALS4 patient cells with control cell lines has revealed a global reduction of cellular R-loop abundance in ALS4 cells (70, 80). In addition, decreased R-loop levels at the promoter of certain genes has also been observed in ALS4 cells and has been linked to the deregulation of a subset of genes (25). Our biochemical data indicate that the ALS4 substitution R2136H does not alter per se SETX catalytic properties leading to an increased R-loop resolving activity (Figure 6F and Supplementary Figure S6E), which could have explained the reduced R-loop levels observed *in cellulo*. However, it has been recently shown that the SETX L389S substitution impairs the interaction of SETX with the zinc finger protein ZPR1, which has been proposed to function as a regulator of SETX activity on R-loops (80). It is therefore possible that the R2136H substitution also interferes with the interaction of SETX with ZPR1 or with ZPR1-mediated regulation of SETX activity.

Alternatively, or additionally, the R2136H substitution might impact on the recently reported function of SETX in stress granules (SGs) (95). SGs are membrane-less organelles that form in the cytoplasm upon exposure to diverse stress conditions and that are highly enriched in RNAs and RNA-binding proteins such as the ALS-associated protein TDP-43 (96). Abnormal SGs have been proposed to nucleate protein inclusions that are typically observed in ALS patients and likely play a role in ALS pathogenesis (97, 98). Among them, TDP-43 aggregates are the most frequently found in all ALS forms, including ALS4 (70, 98). Interestingly, SETX has recently been found to associate with SGs and partake in their disassembly (95). We have observed that the R2136H version of SETX helicase domain has decreased thermostability compared to the wild-type (Figure 6H), which might reflect a higher tendency of the mutant protein to aggregate under certain physicochemical conditions. We therefore speculate that SETX-R2136H might not preserve its proper protein folding and aggregate in SGs naturally forming in motor neurons, thus priming TDP-43 aggregation. The resulting toxic protein inclusions might in turn sequester important cellular factors (*i.e*., mRNAs and RNA-binding proteins), leading to global alterations in transcription and RNA processing that could result in the observed reduction in R-loop levels.

In conclusion, in this study we have provided direct biochemical evidence that SETX is endowed with the ability to resolve R-loops and induce transcription termination and we have generated valuable tools to approach the molecular basis of SETX-associated neurodegenerative diseases. We anticipate that our results will enable better interpretation of the existing *in vivo* data and will possibly orient future studies on the biological function of SETX.

## DATA AVAILABILITY

All primary data are available in this manuscript and supplementary material.

## ACKNOWLEDGEMENT

We thank members of the lab for their support. We thank Olga Jasnovidova (Tallinn University of Technology), Matyas Lanicek, and Martin Matl (Masaryk University) for critically reading the manuscript. We thank Mario Spirek and Jakub Cibulka (Masaryk University) for assistance with BLI assays.

## AUTHOR CONTRIBUTIONS

Experimentation: ZH, OP; Protein purification: VK, OP; *In silico* modelling: RS; Conceptualisation: OP, MS; Funding acquisition: OP, RS, MS; Project administration: OP, RS, MS; Supervision: MS; Writing – original draft: ZH, OP, MS; Writing – review and editing: ZH, VK, OP, RS, MS

