## Supplementary material for "Human senataxin is a bona fide R-loop resolving enzyme and transcription termination factor"

Figures S1-S7.

Table S1-S2.

Supplementary methods.

Supplementary references.

Supplementary figure 1 Hasanova *et* al.

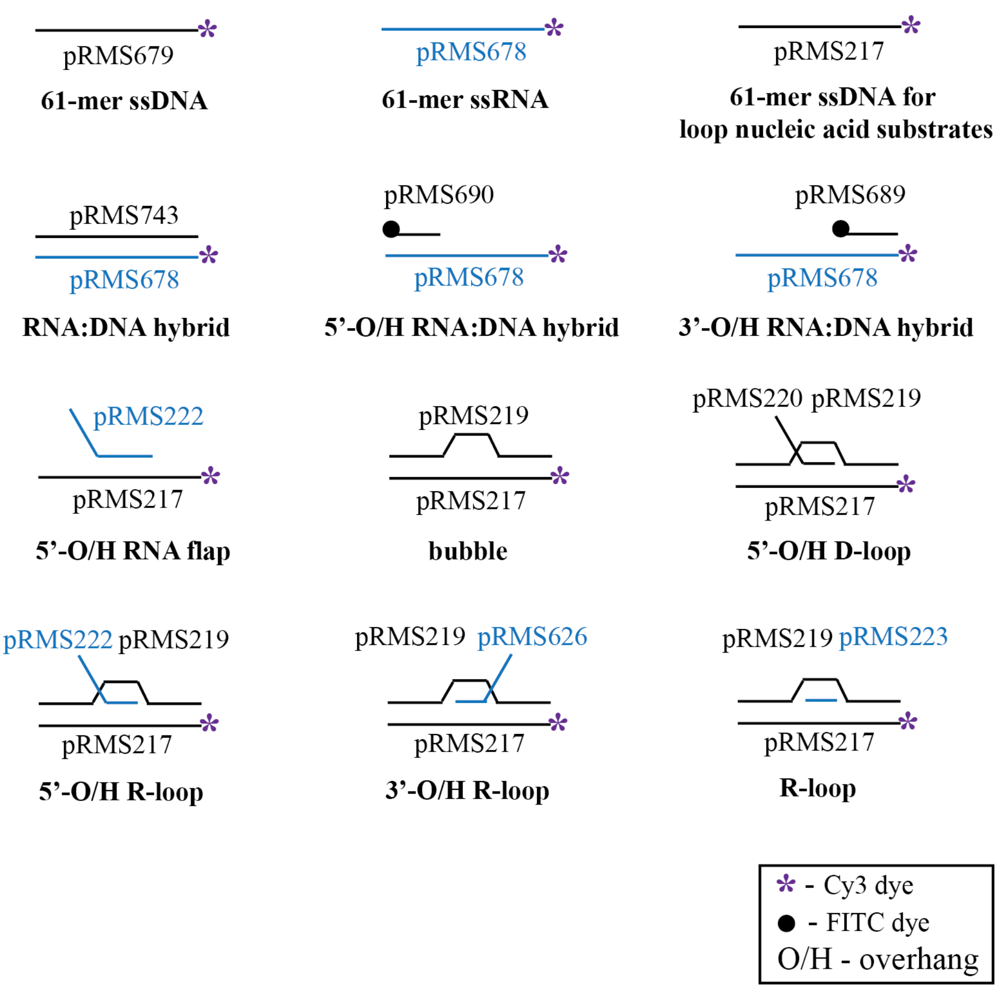


**Figure S1**: **Scheme of used nucleic acid substrates - supplementary figure for Material and Methods.** Depicted are nucleic acid substrates for electrophoretic mobility-shift assays and helicase assays. The names of the constituent oligonucleotides are indicated (see Supplementary table S1 for sequence details). DNA is depicted in black, RNA in blue, Cy3 fluorescent dye with a purple asterisk, and FITC fluorescent dye with a black circle, O/H stands for overhang.

Supplementary figure 2 Hasanova *et* al.

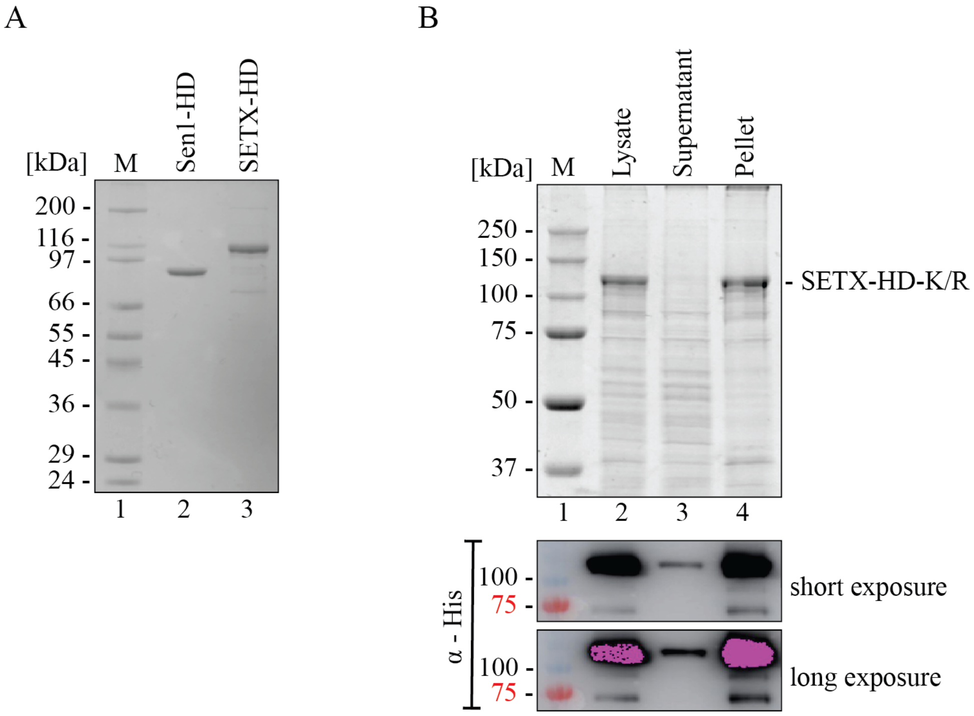


**Figure S2: Supplementary figure to main Figure 1: (A)** SDS-PAGE gel, analysis of purified Sen1-HD and SETX-HD. M: molecular weight marker. (**B)** Solubility test for the mutant version of SETX-HD containing K1969R substitution in the Walker A motif. Top: Coomassie-stained SDS-PAGE gel showing the presence of the SETX-HD-K1969R (K/R) mutant in the cell lysate and in the supernatant and pellet fractions after centrifugation. Bottom: Immunodetection of $(His)_6$-tagged SETX-HD-K/R with an α-His antibody localised SETX-HD-K/R primarily in the pellet fraction (lane 4). M: molecular weight marker.

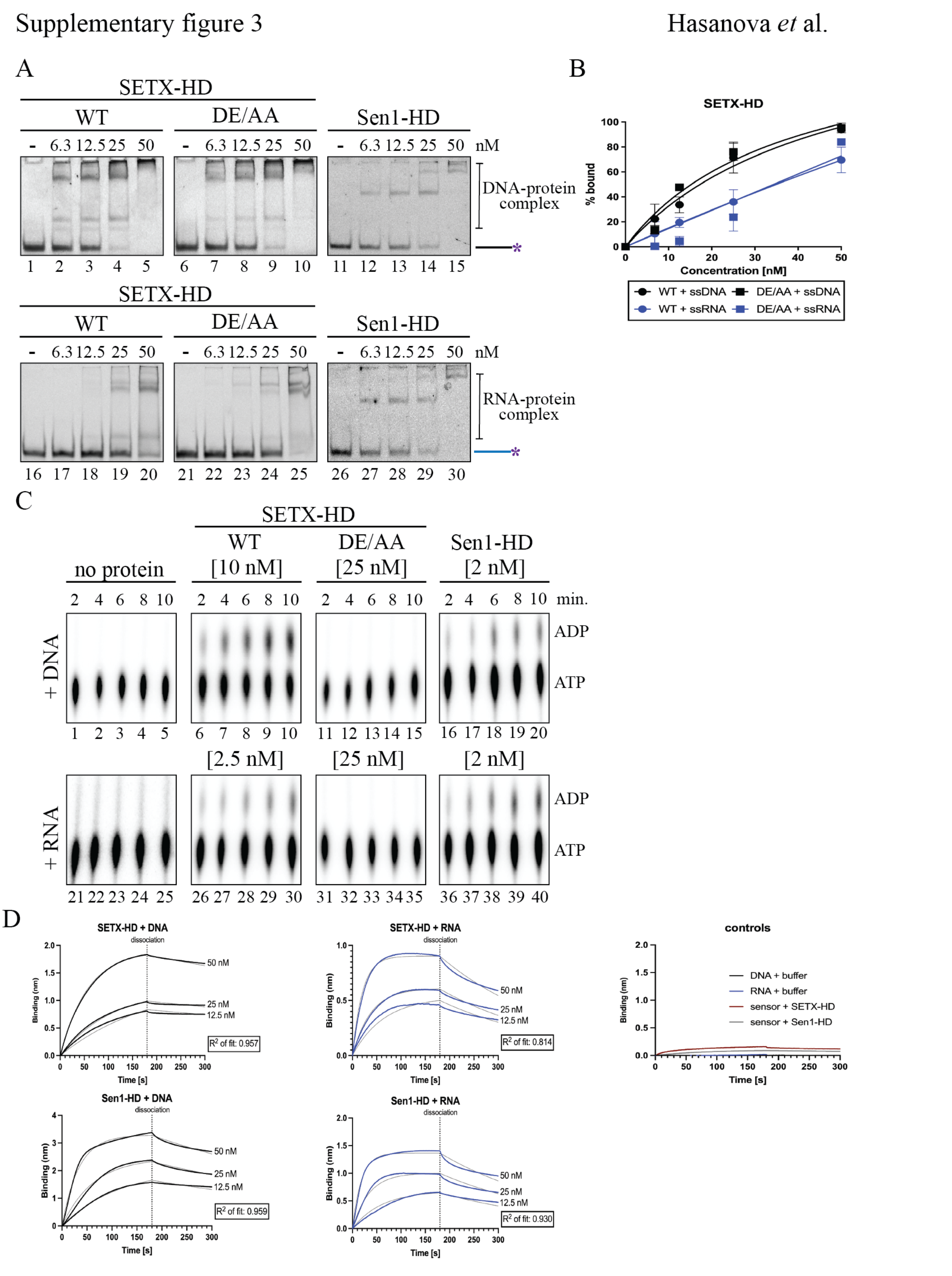


**Figure S3: Supplementary figure to main Figure 2: Biochemical characterisation of SETX-HD-WT and SETX-HD-DE2181, 2182AA (DE/AA). (A)** Electrophoretic mobility-shift assays (EMSA) were performed with the indicated concentrations of SETX-HD-WT, SETX-HD-DE/AA, and Sen1-HD, respectively, in the presence of 10 nM fluorescently labelled ssDNA (pRMS679, lanes 1-15) and ssRNA (pRMS678, lanes 16-30). **(B)** Quantification of nucleic acid binding affinity of SETX-HD-DE/AA compared to SETX-HD-WT from experiments shown in (A). The data points in the graphs represent the mean of 3 independent experiments (for WT + ssDNA: n = 4) and the error bars represent the standard deviation (SD). **(C)** Determination of the ATPase activity of SETX-HD. ATPase assays containing SETX-HD-WT [10 nM or 2.5 nM, as indicated], SETX-HD-DE/AA [25 nM], and Sen1-HD [2 nM], respectively, were performed in the presence of single-stranded DNA (DL3701, lanes 1-20) and RNA (DL3316, lanes 21-40). The position of α[$^{32}$P]-ATP and hydrolysed α[$^{32}$P]-ADP is depicted. **(D)** Sensorgrams from bio-layer interferometry (BLI) assay. Twelve fmoles of biotinylated ssDNA (pRMS739) and ssRNA (pRMS740), respectively, were bound to a SAX biosensor. SETX-HD and Sen1-HD, respectively, were then submerged into the biosensor in three concentrations (12.5, 25, and 50 nM), respectively, to measure protein association (180 s). Protein binding to nucleic acids is represented as the change in light interferometry (in nm). Protein dissociation from nucleic acids was measured in BLI buffer for 120 s. Binding curves (black for DNA and blue for RNA) represent the mean from triplicate measures for each concentration generated in Prism GraphPad 9. Grey lines indicate the non-linear regression fitting curve using the "Association then dissociation" model in global analysis mode. Goodness of fit ($R^2$) is depicted in each graph. Control graph shows association and dissociation of BLI buffer with ssDNA and RNA, respectively, and a sensor without bound nucleic acids with SETX-HD and Sen1-HD, respectively.

Supplementary figure 4 Hasanova *et al.*

A B

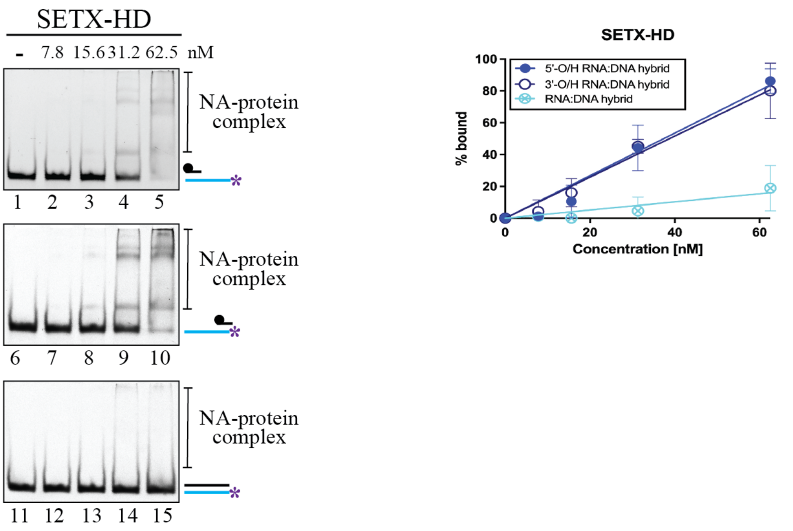


**Figure S4: Supplementary figure to main Figure 3: SETX-HD has similar binding affinity to RNA:DNA hybrids with a single-stranded overhang. (A)** Electrophoretic mobility-shift assays (EMSA) were performed with the indicated concentrations of SETX-HD and fluorescently labelled substrates: 5’-O/H RNA:DNA hybrid (lanes 1-5), 3’-O/H RNA:DNA hybrid (lanes 6-10), and RNA:DNA hybrid with blunt ends (lanes 11-15), all at 10 nM. **(B)** Quantification of the experiments shown in (A). The data points in the graph represent the mean of 3 independent experiments and error bars represent the standard deviation (SD).

Supplementary figure 5 Hasanova *et* al.

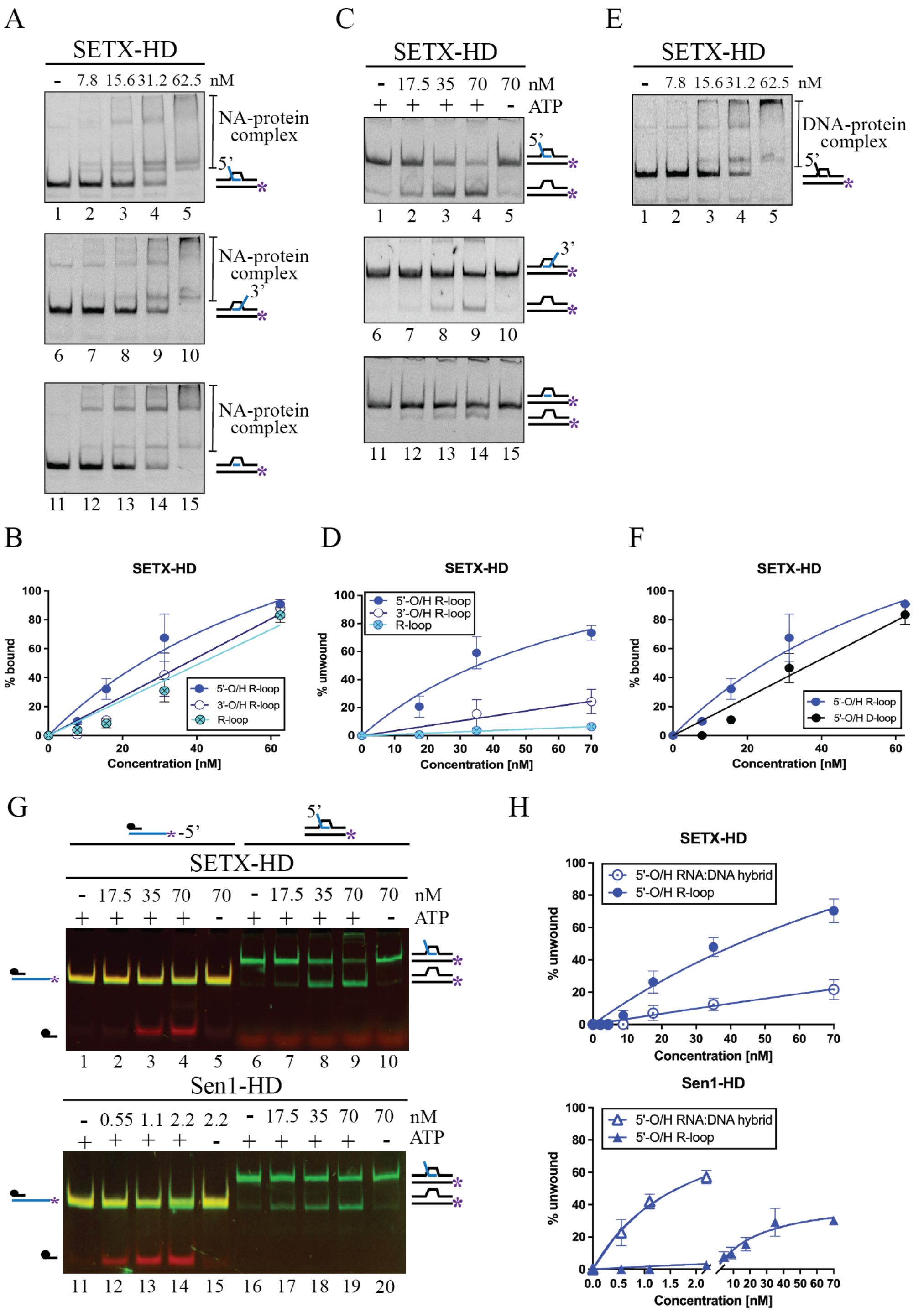


**Figure S5: Supplementary figure to main Figure 4: SETX-HD preferentially unwinds R-loop structures with a 5'-overhang. (A)** SETX-HD binds R-loop structures with similar affinity. Electrophoretic mobility-shift assays (EMSA), which were performed with the indicated amounts of SETX-HD and fluorescently labelled 5'-O/H R- loop (lanes 1-5), 3'-O/H R-loop (lanes 6-10), or R-loop (lanes 11-15), all at 10 nM. **(B)** Quantification of the experiments shown in (A). The data points in the graphs represent the mean of 3 independent experiments and the error bars represent the standard deviation (SD). **(C)** Helicase assays were performed in the presence of the indicated amounts of SETX-HD, and 10 nM fluorescently labelled 5'-O/H R-loop (lanes 1-5), 3'-O/H R-loop (lanes 6-10), or an R-loop with no overhang (lanes 11-15). Where indicated, ATP was omitted. **(D)** Quantification of the experiments shown in (C). The data points in the graphs represent the mean of 3 independent experiments (for WT + 5'-O/H R-loop: n = 5) and the error bars represent the standard deviation (SD). **(E)** Electrophoretic mobility-shift assay (EMSA) was performed with the indicated amounts of SETX-HD and fluorescently labelled 5'-O/H D-loop at 10 nM. **(F)** Quantification of the experiments shown in (E and A - 5'-O/H R-loop - for comparison). The data points in the graphs represent the mean of 3 independent experiments and error bars represent the standard deviation (SD). **(G)** SETX-HD, in contrast to Sen1-HD, preferentially unwinds triplex R-loop structures than duplex RNA:DNA hybrids. Helicase assays were performed in the presence of the indicated amounts of SETX-HD and Sen1-HD, respectively, and 10 nM fluorescently labelled 5'-O/H RNA:DNA hybrid (lanes 1-5 and 11-15) and 5'-O/H R-loop (lanes 6-10 and 16-20). Where indicated, ATP was omitted. Scans are shown as an overlay of FITC (red) and Cy3 (green) signals generated in Multi Gauge software. **(H)** Quantification of the experiments shown in (G). The data points in the graphs represent the mean of 3 independent experiments and error bars represent the standard deviation (SD).

Supplementary figure 6 Hasanova *et* al.

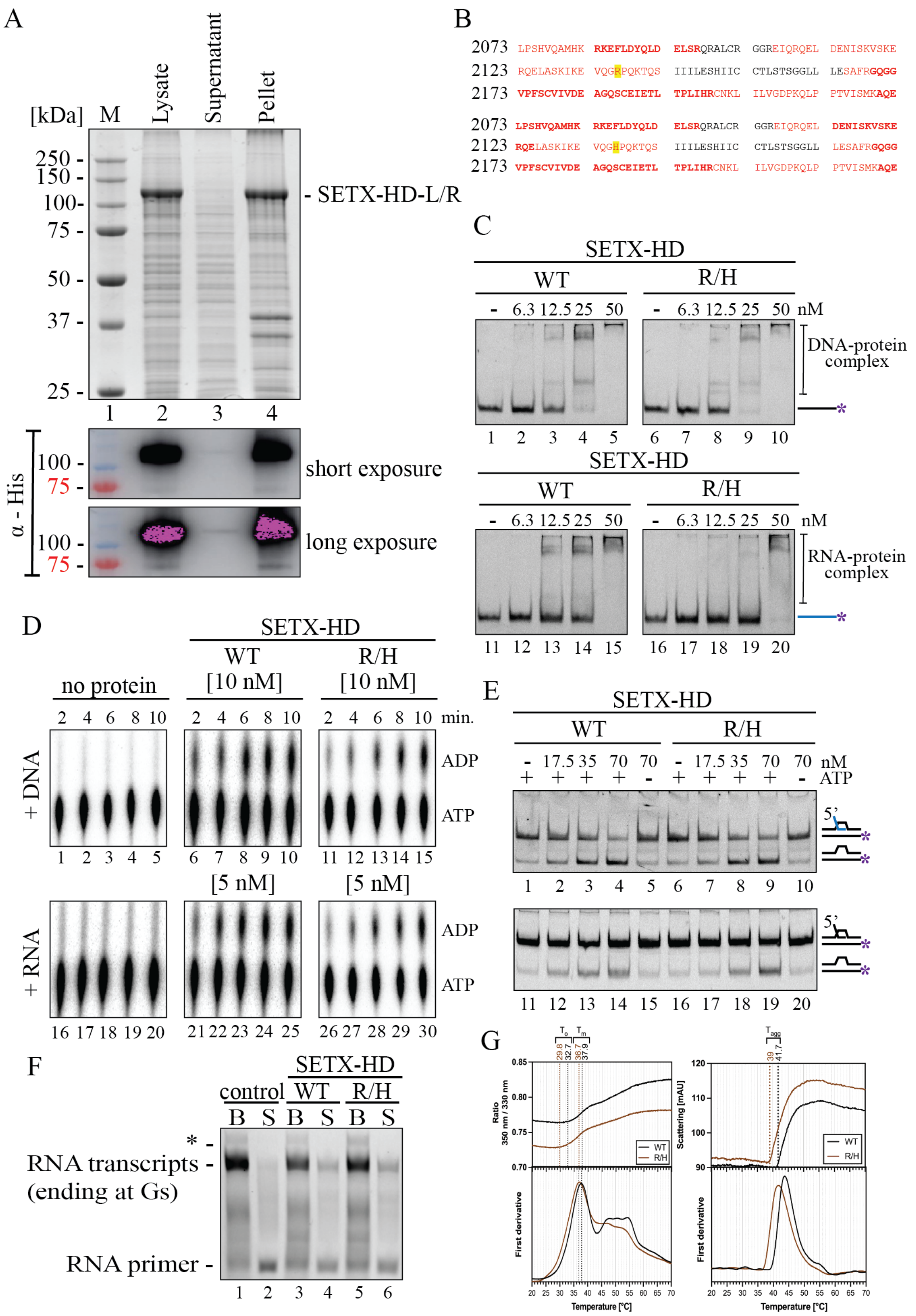


**Figure S6: Supplementary figure to main Figure 6: Purification and characterisation of SETX-HD variants associated with AOA2 and ALS4. (A)** Solubility test for the mutant version of SETX-HD containing the L1976R substitution associated with AOA2. Top: Coomassie-stained SDS-PAGE gel showing the presence of the SETX-HD-L1976R (L/R) mutant in the cell lysate and in the supernatant and pellet fractions after centrifugation. Bottom: Immunodetection of $(His)_6$-tagged SETX-HD-L/R with an α-His antibody localised SETX-HD-L/R primarily in the pellet fraction (lane 4). M: molecular weight marker. **(B)** Mass spectrometry analysis of SETX-HD-WT and SETX-HD-R/H, respectively. Combined amino acid coverage from MALDI-MS/MS and LC-MS/MS. In both proteins the amino acid position of interest (p. 2136) was covered (highlighted in yellow). Covered amino acids are shown in red (bold red depicts significant MASCOT score). **(C)** Electrophoretic mobility-shift assays (EMSA) were performed with the indicated amounts of SETX-HD-WT and SETX-HD-R/H, respectively, in the presence of 10 nM fluorescently labelled ssDNA (pRMS679, lanes 1-10) and ssRNA (pRMS678, lanes 11-20). **(D)** Determination of the ATPase activity of SETX-HD-R/H. ATPase assays containing indicated amounts of SETX-HD-WT and SETX-HD-R/H, respectively, were performed in the presence of single-stranded DNA (DL3701, lanes 1-15) and RNA (DL3316, lanes 16-30). The position of α[$^{32}$P]-ATP and hydrolysed α[$^{32}$P]-ADP is depicted. **(E)** SETX-HD-R/H unwinds 5'-O/H R-loop and 5'-O/H D-loop structures, respectively, to a similar extent as SETX-HD-WT. Helicase assays were performed in the presence of the indicated concentrations of SETX-HD-WT and SETX-HD-R/H (17.5, 35, and 70 nM), respectively, and 10 nM fluorescently labelled 5'-O/H R-loop (upper panel) or 5'- O/H D-loop (lower panel) in the presence of ATP. Lanes 5, 10, 15, and 20 represent control experiments with no ATP. **(F)** SETX- HD-R/H induces transcription termination *in vitro* to a similar extent as the wild-type. *In vitro* transcription termination assays performed with mammalian RNAPII as in Figure 5. Asterisk denotes transcripts associated with RNAPIIs that over-read the G-stretch due to small GTP contaminations. **(G)** Representative nanoDSF thermogram for SETX-HD-WT (in black) and SETX-HD-R/H (in brown). Upper left panel shows fluorescence intensity as a 350/330 nm ratio indicating protein unfolding and the upper right panel shows light scattering (back reflected light intensity passing through the sample) indicating the onset of protein aggregation. Corresponding first derivatives are plotted in the bottom panel. Temperatures of onset of protein unfolding ($T_o$), melting temperature ($T_m$), and onset of protein aggregation ($T_a$) are shown as vertical lines.

Supplementary figure 7 Hasanova *et al.*

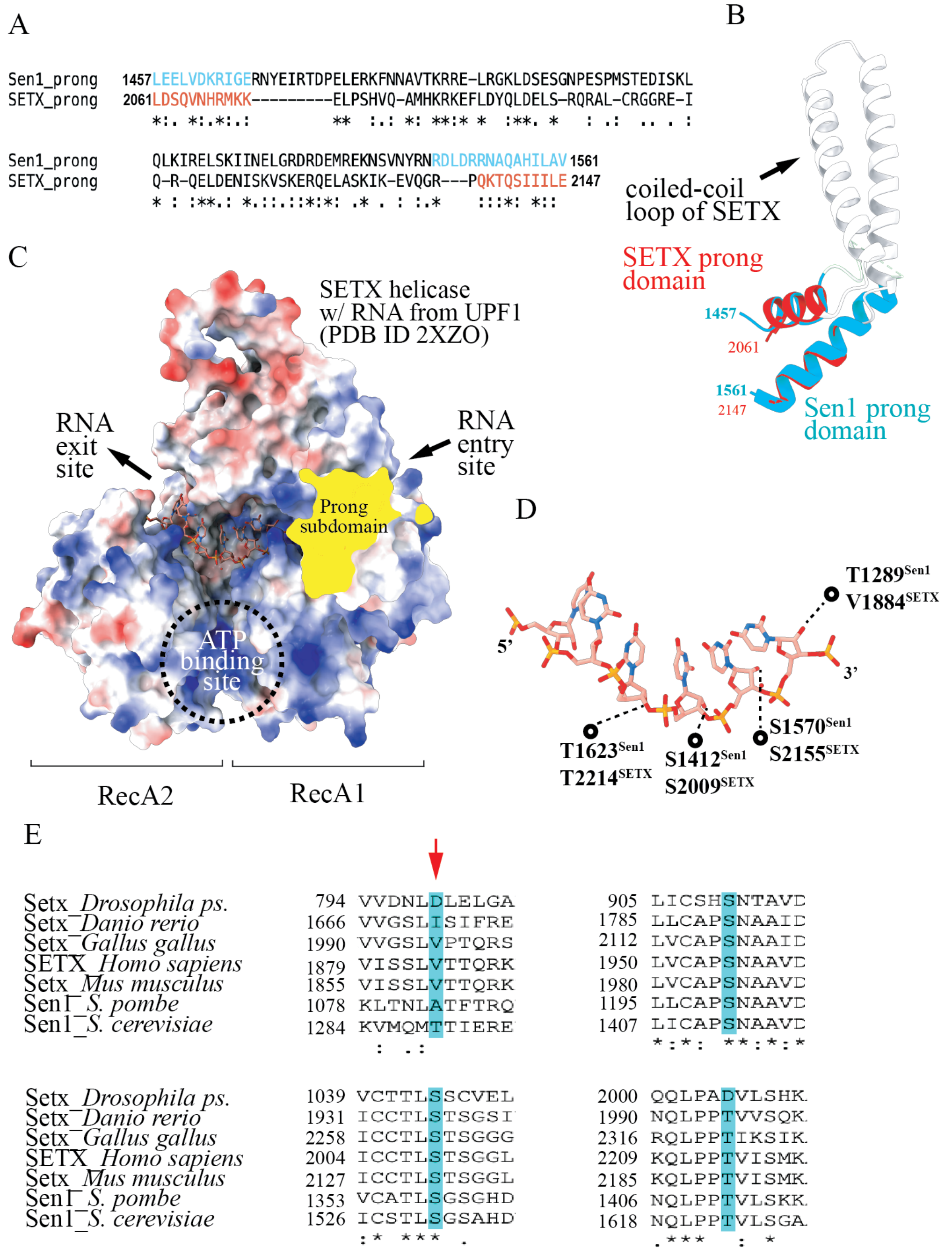


**Figure S7: Comparison of the prong domain from Sen1 and SETX and nucleic acids binding in the helicase channel. (A)** Structure-guided amino acid alignment of the prong domain from Sen1 and SETX. **(B)** Overlay of the prong domain from the crystal structure of Sen1 (PDB ID 5MZN, in blue) and from the AlphaFold2 model of SETX (in red). **(C)** SETX model (shown as electrostatic surface) with RNA from UPF1-RNA complex (2XZO, (1)). RNA was placed in the helicase channel based on the overlay of the structures that are highly similar. **(D)** Recognition of the RNA sugar moieties (shown as sticks) in the helicase channel of SETX and Sen1. **(E)** Multiple sequence alignment of RNA sugar binding sites in Sen1/SETX in various organisms. The area (+/- 5 amino acids) around the RNA sugar recognition sites (highlighted in blue) of the helicase channel (presented in (D)) for *Drosophila pseudoobscura*, *Danio rerio*, *Gallus gallus*, *Homo sapiens*, *Mus musculus*, *Schizosaccharomyces pombe* and *Saccharomyces cerevisiae*. The red arrow shows the position of the highly variable RNA recognition site (in humans V1884). Multiple sequence alignment was done using the ClustalW online tool.

**Supplementary table S1: List of oligonucleotides used in this work.**

| Name | Sequence 5'-3' | Use |
|---|---|---|
| pRMS101 | ATCCACGGACCACCTGGAACCGGAAGGTCT AAGACCATCGTCGGACTG | Forward primer for site-directed mutagenesis to introduce K1969R mutation. |
| pRMS102 | CAGTCCGACGATGGTCTTAGACCTTCCGGTT CCAGGTGGTCCGTGGAT | Reverse primer for site-directed mutagenesis to introduce K1969R mutation. |
| pRMS103 | CCCTTCTCTTGCGTGATCGTGGCCGCAGCCG GACAGTCTTGCGAAATCGAAACTC | Forward primer for site-directed mutagenesis to introduce DE2181, 2182AA mutation. |
| pRMS104 | GAGTTTCGATTTCGCAAGACTGTCCGGCTGC GGCCACGATCACGCAAGAGAAGGG | Reverse primer for site-directed mutagenesis to introduce DE2181, 2182AA mutation. |
| pRMS114 | ACCTCCTGCCTCTCTGTGC | Sequencing of SETX |
| pRMS115 | GGACTTCAACGAGGACCAAA | Sequencing of SETX |
| pRMS116 | CTGCTGCACCTTGTCTACCA | Sequencing of SETX |
| pRMS117 | AGTCGACACCGTTGATGCTT | Sequencing of SETX |
| pRMS217 | GACGCTGCCGAATTCTACCAGTGCCTTGCTA GGACATCTTTGCCCACCTGCAGGTTCACCC | 61-mer DNA primer fluorescently labelled on 5' end with Cy3 used to create R-loop/D-loop/bubble substrates for EMSAs and helicase assays. |
| pRMS219 | GGGTGAACCTGCAGGTGGGCGGCTGCTCAT CGTAGGTTAGTTGGTAGAATTCGGCAGCGTC | 61-mer DNA primer used to create R-loop/D-loop/bubble substrates for EMSAs and helicase assays. |
| pRMS220 | TAAGAGCAAGATGTTCTATAAAAGATGTCCT AGCAAGGCAC | 41-mer DNA primer used to create a 5'-overhang D-loop substrate for EMSAs and helicase assays. |
| pRMS222 | UAAGAGCAAGAUGUUCUAUAAAAGAUGUC CUAGCAAGGCAC | 41-mer RNA primer used to create a 5'-overhang R-loop and RNA:DNA hybrid substrate for EMSAs and helicase assays. |
| pRMS223 | AAAGAUGUCCUAGCAAGGCAC | 21-mer RNA primer used to create an R-loop substrate for EMSAs and helicase assays. |
| pRMS323 | TTATCCACTTCCAATGTTATTATTAGTGATG GTGATGGTGATG | Reverse primer for generating a 1674-2677 amino acid fragment of SETX for cloning into 438A plasmid. |
| pRMS590 | TTTAAGAAGGAGATATAGTTCATGGTCCTGA ACACCTTCG | Forward primer for generating a 1674-2677 amino acid fragment of SETX for cloning into 2BcT plasmid. |
| pRMS591 | GGATTGGAAGTAGAGGTTCTCGAGCAACTT ACGCTTCTTTG | Reverse primer for generating a 1674-2677 amino acid fragment of SETX for cloning into 2BcT plasmid. |
| pRMS592 | TACTTCCAATCCAATCGATGGTCCTGAACAC CTTCG | Forward primer for generating a 1674-2677 amino acid fragment of SETX for cloning into 438A plasmid. |
| pRMS607 | GGAAAGTCTAAGACCATCGTCGGACGGCTG TACAGGCTGCTCACC | Forward primer for site-directed mutagenesis to introduce a L1976R mutation. |
| pRMS608 | GGTGAGCAGCCTGTACAGCCGTCCGACGAT GGTCTTAGACTTTCC | Reverse primer for site-directed mutagenesis to introduce a L1976R mutation. |
| pRMS609 | TCCAAGATCAAAGAGGTCCAGGGA<u>CAC</u>CCT CAGAAAACCCAGTCT | Forward primer for site-directed mutagenesis to introduce a R2136H mutation. |

| Name | Sequence 5'-3' | Use |
|---|---|---|
| pRMS610 | AGACTGGGTTTTCTGAGGGTGTCCCTGGACCTCTTTGATCTTGGA | Reverse primer for site-directed mutagenesis to introduce a R2136H mutation. |
| pRMS626 | AAAGAUGUCCUAGCAAGGCACUAAGUCCUAGAUGUUCUAUA | 41-mer RNA primer used to create a 3'-overhang R-loop substrate for EMSAs and helicase assays. |
| pRMS678 | UUGUCUUCUCAUAAAUUUAAUCCCCGUACGCUUAUACUCCUUUAACUACCCAGUCUUCCCG | 61-mer RNA primer fluorescently labelled on 5' end with Cy3 used as a single-stranded RNA for EMSAs and to create RNA:DNA hybrid substrates for EMSAs and helicase assays. |
| pRMS679 | TTGTCTTCTCATAAATTTAATCCCCGTACGCTTATACTCCTTTAACTACCCAGTCTTCCCG | 61-mer DNA primer fluorescently labelled on 5' end with Cy3 used as a single-stranded DNA for EMSAs. |
| pRMS680 | TTGTCTTCTCATAAATTTAATCCCCGTACGCTTATACTCCTTTAACTACCCAGTCTTCCCG | 61-mer DNA primer used as a competitor in helicase assays with a RNA:DNA hybrid. |
| pRMS681 | TTAAATTTATGAGAAGACAA | 20-mer DNA primer used as a competitor in helicase assays with a 3'-overhang RNA:DNA hybrid. |
| pRMS682 | CGGGAAGACTGGGTAGTTAA | 20-mer DNA primer used as a competitor in helicase assays with a 5'-overhang RNA:DNA hybrid. |
| pRMS689 | TTAAATTTATGAGAAGACAA | 20-mer DNA primer fluorescently labelled on 5' end with FITC used to create a 3'-overhang RNA:DNA hybrid substrate for EMSAs and helicase assays. |
| pRMS690 | CGGGAAGACTGGGTAGTTAA | 20-mer DNA primer fluorescently labelled on 5' end with FITC used to create a 5'-overhang RNA:DNA hybrid substrate for EMSAs and helicase assays. |
| pRMS739 | TTGTCTTCTCATAAATTTAATCCCCGTACGCTTATACTCCT | 41-mer 5' biotin tagged DNA primer for BLI assay |
| pRMS740 | UUGUCUUCUCAUAAAUUUAAUCCCCGUACGCUUAUACUCCU | 41-mer 5' biotin tagged RNA primer for BLI assay |
| pRMS743 | CGGGAAGACTGGGTAGTTAAAGGAGTATAAGCGTACGGGGATTAAATTTATGAGAAGACAA | 61-mer DNA primer used to create a blunt ended RNA:DNA hybrid for EMSAs and helicase assays. |
| DL3701 | TTCATTTCAGACCAGCACCCACTCACTACAACTCACGACCAGGC | Primer used to stimulate nucleic-acid dependent SETX/Sen1 ATPase activity. |
| DL3316 | UUCAUUUCAGACCAGCACCCACUCACUACAACUCACGACCAGGC | Primer used to stimulate nucleic-acid dependent SETX/Sen1 ATPase activity. |
| DL2492 | UGCAUUUCGACCAGGC | 5' FAM labelled RNA primer for performing IVTT assays on immobilized templates. |
| DL3352 | CTAGAGGAAACAAACTATAGGAAACGACCAGGCCCTCAACATCTCTCACCCATCTCCACACGGGGGTTACCCGGCCTGCA | Non-template strand for IVTT assays. |
| DL3353 | GGCCGGGTAACCCCCGTGTGGAGATGGGTGAGAGATGTTGAGGGCCTGGTCGTTTCCTATAGTTTGTTTCCT | Template strand for IVTT assays. |

**Supplementary table S2: List of proteins identified by MS analysis in a typical SETX-HD purification.**

SETX-HD wild-type

| NCBIprot entry | Protein | Score | Organism | Sequence coverage [%] |
|---|---|---|---|---|
| BAG10365.1 | senataxin (fragment) | 10135 | *Homo sapiens* | 82 |
| XP_026726793.1 | ras GTPase-activating protein-binding protein 2 | 1141 | *Trichoplusia ni* | 45 |
| XP_026736716.1 | cleavage and polyadenylation specificity factor subunit CG7185 | 543 | *Trichoplusia ni* | 19 |
| XP_026740726.1 | protein lingerer-like | 511 | *Trichoplusia ni* | 23 |
| XP_026743046.1 | AMP deaminase 2-like isoform X6 | 503 | *Trichoplusia ni* | 34 |
| XP_026748099.1 | cytoplasmic FMR1-interacting protein | 376 | *Trichoplusia ni* | 18 |
| XP_026739518.1 | transcription elongation factor SPT6 | 365 | *Trichoplusia ni* | 19 |
| XP_026733954.1 | protein LSM14 homolog A | 323 | *Trichoplusia ni* | 22 |
| XP_026747555.1 | la-related protein 1-like | 292 | *Trichoplusia ni* | 28 |
| XP_026747903.1 | cleavage and polyadenylation specificity factor subunit 5 | 265 | *Trichoplusia ni* | 35 |
| XP_026730705.1 | ataxin-2-like protein, partial | 194 | *Trichoplusia ni* | 21 |

SETX-HD R2136H mutant

| NCBIprot entry | Protein | Score | Organism | Sequence coverage [%] |
|---|---|---|---|---|
| BAG10365.1 | senataxin (fragment) | 11213 | *Homo sapiens* | 82 |
| XP_026743046.1 | AMP deaminase 2-like isoform X6 | 962 | *Trichoplusia ni* | 48 |
| XP_026736716.1 | cleavage and polyadenylation specificity factor subunit CG7185 | 672 | *Trichoplusia ni* | 18 |
| XP_026726793.1 | ras GTPase-activating protein-binding protein 2 | 613 | *Trichoplusia ni* | 38 |
| XP_026739518.1 | transcription elongation factor SPT6 | 474 | *Trichoplusia ni* | 25 |
| XP_026747555.1 | la-related protein 1-like | 449 | *Trichoplusia ni* | 32 |
| XP_026732337.1 | T-complex protein 1 subunit delta | 417 | *Trichoplusia ni* | 45 |
| XP_026740726.1 | protein lingerer-like | 319 | *Trichoplusia ni* | 13 |
| AAB06239.1 | HSC70 | 259 | *Trichoplusia ni* | 39 |
| XP_026747903.1 | cleavage and polyadenylation specificity factor subunit 5 | 225 | *Trichoplusia ni* | 34 |
| XP_026730705.1 | ataxin-2-like protein, partial | 194 | *Trichoplusia ni* | 21 |

### Supplementary methods

#### Protein identification from 1D gel bands by LC-MS/MS

The proteins were subjected to incubation with trypsin or pepsin for 1 hour at 37 °C. The resulting peptides were analyzed by LC-MS/MS using RSLCnano system (Thermo Fisher Scientific) connected to Qq-Time-Of-Flight mass spectrometer Impact II (Bruker Daltonics). MS/MS data were searched against a custom database of relevant protein sequences in combination with cRAP contaminant database, and against NCBIprot database (downloaded on $6^{th}$ February 2021) without taxonomy restrictions, using an in-house Mascot search engine (Matrixscience; version 2.6).

#### Solubility test and immunodetection of SETX-HD variants by western blotting

Hi5 cell pellets with expressed SETX-HD variants were lysed in lysis buffer (50 mM Tris-HCl, pH 8; 0.5 M NaCl; 10 % (v/v) glycerol; 1 mM DTT; 0.4 % (v/v) Triton-X; 10 mM imidazole) containing protease inhibitors (0.66 µg/ml pepstatin, 5 µg/ml benzamidine, 4.75 µg/ml leupeptin, 2 µg/ml aprotinin) and 25 U benzonase per ml of lysate. The lysate was cleared by centrifugation to obtain soluble proteins. The individual fractions – lysate, supernatant, and pellet after centrifugation were separated on SDS-PAGE and transferred on a PVDF membrane (AppliChem). Monoclonal mouse anti-His antibodies (SAB1305538) and goat anti-mouse IgG conjugated with HRP (A0168) were purchased from Sigma-Merck.
